# UFD1 Recognition of Initiator and Proximal Ubiquitin Drives p97-Mediated Substrate Unfolding enhanced by FAF1, FAF2, and UBXD7

**DOI:** 10.64898/2026.03.13.711538

**Authors:** Shunki Miyauchi, Ami Hirano, Nanako Yasumoto, Yuiha Goto, Yoshino Akizuki, Fumika Koyano, Tomoya Hino, Shingo Nagano, Noriyuki Matsuda, Fumiaki Ohtake, Gosuke Hayashi, Yusuke Sato

**Affiliations:** Department of Engineering, Graduate School of Sustainability Science, Tottori University, Tottori 680-8552, Japan; Faculty of Engineering, Tottori University, Tottori 680-8552, Japan; Department of Biomolecular Engineering and Institute of Nano-Life-Systems, Institute of innovation for Future Society, Nagoya University, Nagoya 464-8603, Japan; Laboratory of Protein Degradation, Institute for Advanced Life Sciences, Hoshi University, Tokyo 142-8501, Japan; Graduate School of Pharmacy and Pharmaceutical Sciences, Hoshi University, Tokyo 142-8501, Japan; Department of Biomolecular Pathogenesis, Institute of Integrated Research, Institute of Science Tokyo, Tokyo 113-8510, Tokyo, Japan; Department of Chemistry and Biotechnology, Graduate School of Engineering, Tottori University, Tottori 680-8552, Japan; Center for Research on Green Sustainable Chemistry, Tottori University, Tottori 680-8552, Japan; Chromosome Engineering Research Center, Tottori University, Tottori 683-8503, Japan

**Author notes:** Correspondence should be addressed to Y.S.

## Abstract

The AAA-ATPase p97 and its primary adaptor UFD1–NPL4 (UN) unfold ubiquitinated substrates for their proteasomal degradation. Although cryo-EM has shown that substrate processing begins with unfolding of the initiator ubiquitin and its capture by NPL4, the specific contribution of UFD1 to K48-linked ubiquitin chain engagement has not been fully defined. Here we report a 1.31 Å crystal structure of the human UFD1-UT3 bound to a K48-linked di-ubiquitin mimic, revealing that UFD1-UT3 engages the unfolded initiator ubiquitin via its Nc subdomain and the proximal ubiquitin via its Nn subdomain. AlphaFold3-guided analyses further demonstrate that accessory adaptors FAF1, FAF2, and UBXD7 enhance p97–UN activity through divergent strategies. FAF1 and FAF2 scaffold UFD1-UT3 near the NPL4-tower, whereas UBXD7 stabilizes transient interactions between UFD1 and the ubiquitin chain. Our results provide a comprehensive molecular model for the initiation of p97-mediated substrate unfolding.

## Introduction

Protein homeostasis relies on poly-ubiquitin (Ub) chains linked through Lys48 (K48-chain), which serve as the canonical signal for proteasomal degradation^1–3^. Positioned upstream of the Ub–proteasome system (UPS)^4^, the AAA-ATPase p97 (also known as VCP in mammals or Cdc48 in yeast) provides the mechanical force for substrate extraction^5–7^. Driven by the energy of ATP hydrolysis, p97 extracts and unfolds ubiquitinated proteins from various cellular environments, including the endoplasmic reticulum (ERAD), mitochondria, and chromatin^8^. Over 30 distinct adaptors recruit the p97 hexamer to specific sites and modulate its activity^9^. Among these, the UFD1–NPL4 (UN) heterodimer acts as the primary adaptor recognizing K48-chains for protein unfolding^4,10–13^.

High-resolution cryo-EM studies of the yeast and human p97–UN have shown that this substrate-processing mechanism is evolutionarily conserved^13–16^. The reaction initiates when the first Ub molecule in a chain, termed the initiator Ub (Ub^init^), is unfolded and captured by NPL4 in an ATP-independent manner^13^. This unfolded polypeptide is then threaded through the central pore of the p97 hexameric ring, in which sequential ATP hydrolysis drives its translocation. Despite these insights, the structural mechanism by which the complex achieves the high-affinity engagement required to initiate unfolding of a folded Ub moiety remains a fundamental question. NPL4 is the primary receptor for unfolded Ub^13^, but UFD1 is the key determinant for the initial selection of K48 chains^15^. Although the UFD1-UT3 domain interacts with K48-chain^17,18^, the molecular details have remained unresolved due to its characteristically weak affinity of UFD1 for native, folded K48-chains. Efficient initiation may require UFD1 to interact with the proximal Ub (Ub^prox^) adjacent to Ub^init^ to ensure proper substrate orientation and residency time^15^. However, the intrinsic flexibility of UFD1-UT3 has made it exceptionally challenging to identify the structural determinants governing this critical engagement^13,19^.

Accessory adaptors such as UBXD7, FAF1, and FAF2 further refine unfolding efficiency and regulatory precision^9,20^. These adaptors share a modular UBA-UAS-UBX architecture but govern diverse processes^21,22^. FAF1 and UBXD7 mediate the degradation of substrates in ERAD and chromatin pathways^21,23–26^, while FAF2 promotes peroxisomal membrane protein turnover^7,27^. Recent studies indicate that these adaptors accelerate substrate unfolding, particularly for short Ub chains^21,22^. Recent work has reported that FAF1 and FAF2 position the UFD1-UT3 domain for efficient loading^22^. In contrast, although UBA or UAS domains are essential for UBXD7-mediated enhancement, the underlying molecular mechanism remains elusive^21,22^. Thus, the physical nature of this regulation and what structural strategies these diverse adaptors employ to enhance the unfolding activity of p97–UN remain to be elucidated.

Here we report the crystal structure of the human UFD1-UT3 domain in complex with a K48-linked di-Ub mimic at a high resolution of 1.31 Å. This structure provides definitive evidence that UFD1-UT3 coordinates a K48-linked di-Ub (K48-Ub_2_) in which the distal Ub is stabilized in an unfolded state. The use of AlphaFold3 (AF3) structural predictions was crucial in this research^28^, as it enabled the rapid identification of binding partners and accelerated the elucidation of the underlying regulatory mechanisms. Our crystal structure reveals that the Nc subdomain groove of UFD1-UT3 specifically coordinates the C-terminal segment of an unfolded Ub^init^, whereas the Nn subdomain recognizes the Ub^prox^. This structural arrangement provides the first atomic-level evidence of how UFD1 establishes its unique specificity for K48-chains by stabilizing the Ub^init^ in an unfolded conformation, thereby preventing spontaneous refolding. Furthermore, we reveal that FAF1, FAF2, and UBXD7 enhance p97–UN activity through divergent structural strategies that converge on the UFD1-UT3 module. FAF1 and FAF2 employ a rigid helical motif to scaffold UFD1-UT3 into a functional orientation near the NPL4-tower. In contrast, our findings suggest that UBXD7 promotes the reaction by stabilizing the transient interaction between UFD1-UT3 and Ub^prox^. These results define the structural basis of UFD1-mediated initiation of unfolding and identify the Ub^init^-binding groove as a promising target for designing next-generation, UPS-specific p97 inhibitors.

## Results

### The UFD1-UT3 domain engages K48-linked chains possessing an unfolded initiator ubiquitin

To elucidate the function of the UT3 domain of UFD1 (UFD1-UT3) during the unfolding process, we performed structure predictions using AF3^28^. While initial inputs using full-length UFD1 yielded low-confidence results, we input only the interacting domains^29^, one chain of UFD1-UT3 and two chains of Ub. Notably, despite the absence of predefined distance restraints, the prediction yielded a structure mimicking the UFD1–UT3–K48Ub_2_ complex, in which Gly76 of the distal Ub (Ub^dist^) was positioned near the Lys48 of the Ub^prox^ at a distance compatible with isopeptide bond formation (Extended Data Fig. 1a). In this model, Ub^prox^ stably associated with the Nn subdomain, whereas Ub^dist^ displayed high flexibility at the Nc subdomain, except for its C-terminal tail (residues 73-76). The five structures outputted by AF3 showed significant deviations in the Ub^dist^ structures. Notably, the Ub^prox^-binding surface identified on the Nn subdomain aligns with previously reported NMR analyses characterizing the interaction between UFD1-UT3 and K48 chains^18^.

Next, given that the C-terminus of Ub^init^ (residues 51–76) remained unresolved in prior cryo-EM studies^13,19^, we hypothesized that this segment interacts with UFD1-UT3 upon unfolding. We performed AF3 predictions using a truncated Ub fragment (residues 51–76) to represent the unfolded state. The resulting structural model, hereafter referred to as the UFD1-UT3–K48-Ub ^init^ complex, exhibited substantially enhanced structural confidence, as evidenced by markedly improved PAE maps compared to the folded K48-Ub_2_ complex (Extended Data Fig. 1a, b). In this complex, the Ub^init^ C-terminus (residues 68–76) was deeply accommodated within a previously uncharacterized groove of the UFD1-UT3 Nc subdomain.

To validate the predicted structural model, we chemically synthesized K48-Ub ^init^, in which the C-terminal segment of Ub^init^ (residues 65–76) is linked to Lys48 of Ub^prox^ via an isopeptide bond (Extended Data Fig. 2a). We determined the crystal structure of the UFD1-UT3 in complex with K48-Ub ^init^ at 1.31 Å resolution (Fig. 1a and Table 1). The isopeptide bond within the chemically synthesized K48-Ub ^init^ was well ordered, with unambiguous electron density observed for the Lys48–Gly76 linkage (Extended Data Fig. 2b). In the native fold, residues 64-72 of Ub are embedded within an intramolecular β-sheet, thereby prevents their engagement with the Nc subdomain groove (Fig. 1b). However, the unfolding of Ub^init^ exposes this segment to dock into the Nc subdomain groove and establish an intermolecular β-sheet with UFD1-UT3.

**Figure 1.**
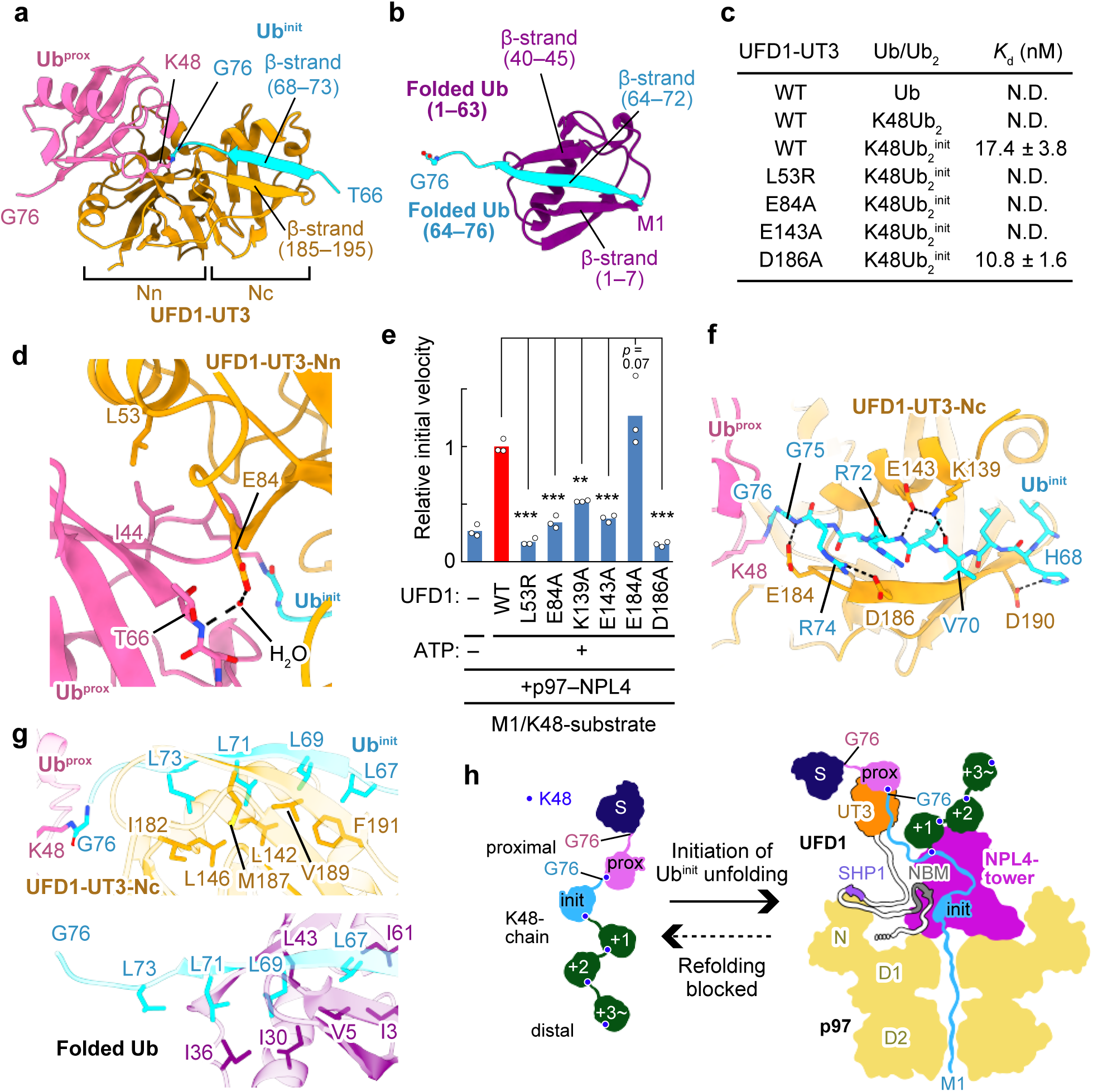
UFD1–UT3 binds to the unfolded initiator ubiquitin and the proximal ubiquitin. **a,** Overall structure of the UFD1-UT3–K48-Ub ^init^ complex. UFD1-UT3, Ub^prox^, and Ub^init^ are colored orange, pink, and cyan, respectively. **b,** Overall structure of Ub (PDB ID:1UBQ)^48^. Residues 1-63 and 64-76 are colored dark purple and cyan, respectively. **c,** Binding affinity of UFD1-UT3 for Ub, K48-Ub_2_, or K48-Ub ^init^. Data are presented as mean ± standard deviation; n = 3 independent experiments. N.D., Not determined. **d,** Close-up view of the interactions between UFD1-UT3 and Ub^prox^. Hydrogen bonds are shown as black dotted lines. **e,** Relative initial velocity of the unfolding of M1/K48-substrate by p97–UN containing WT or mutant UFD1. Individual data points are shown as white dots, and bars represent the mean (n = 3 independent experiments). Relative initial velocities were normalized to the mean of the p97–UN complex in the presence of ATP (red bar). Statistical significance was determined using one-way ANOVA followed by Dunnett’s *post hoc* tests. \*\*\**p* < 0.001; \*\**p* < 0.01. Source data are provided in Extended Data Fig. 4. **f,** Close-up view of the hydrophilic interactions between UFD1-UT3 and Ub^init^. Hydrogen bonds and ionic bonds are shown as black dotted lines. **g,** Structural organization of the Leu cluster at the C terminus of Ub. The top panel shows the conformation of the UFD1-UT3–Ub^init^ interface within the UFD1-UT3–K48Ub_2_^init^ complex. The bottom panel displays the corresponding Leu cluster in folded Ub (PDB ID:1UBQ)^48^ for structural comparison. **h,** Schematic model for the initiation of unfolding of K48-chain modified substrates by the p97–UN complex. S: substrate; prox: Ubprox; init: Ub^init^. Ub distal to the Ub^init^ are numbered +1, +2, and +3 in order of proximity to Ub^init^. Blue dots indicate K48 residues.

**Table 1.**
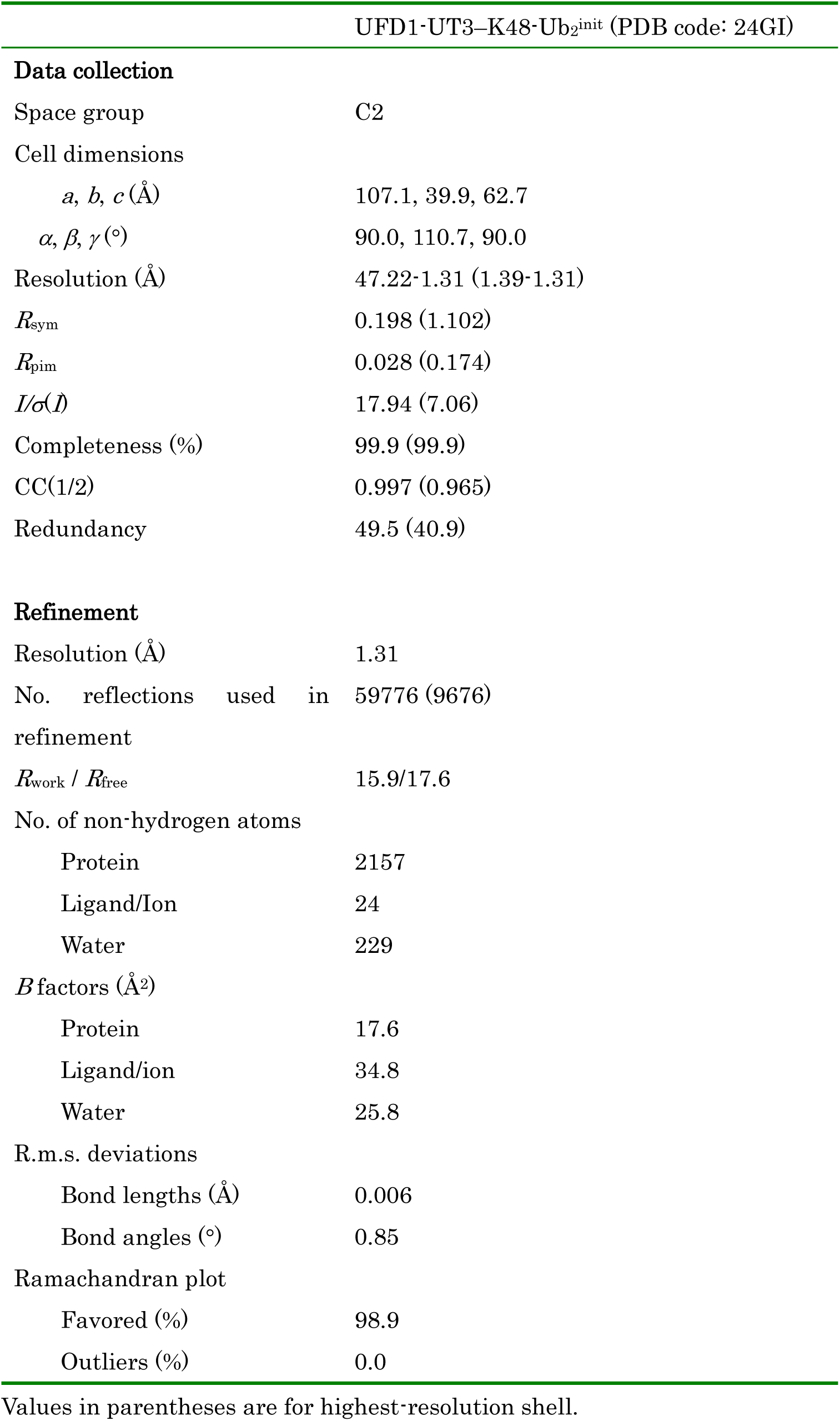
Data collection and refinement statistics.

Bio-layer interferometry (BLI)^30^ further revealed that while UFD1-UT3 binds to mono-Ub and folded K48-Ub_2_ negligibly, K48-Ub ^init^ displays robust affinity (with a *K*_d_ of 17.4 nM) (Fig. 1c and Extended Data Fig. 2c, d). These results align with earlier NMR and HDX-MS reports, revealing that the Nc subdomain does not interact with folded K48-chains but becomes a critical binding site as the chain undergoes unfolding^13,18^. This high-affinity interaction provides a molecular basis for selective substrate recognition during p97-mediated processing.

### Proximal and Initiator Ubiquitin recognition by the UFD1-UT3 domain is essential for p97-dependent unfolding

In the UFD1-UT3–K48-Ub_2_ ^init^ structure, Ile44 of Ub^prox^ interacts with Leu53 of UFD1, while the main-chain NH group of Thr66 in Ub^prox^ is water-bridged to Glu84 of UFD1 (Fig. 1d). To assess these roles, we introduced L53R and E84A mutations into UFD1 and analyzed the p97–UN unfolding activity and binding affinity toward K48-Ub ^init^. For unfolding assays, we prepared mEos3.2 modified with a K48-chain or an M1/K48-branched chain (designated as the K48-substrate or M1/K48-substrate, respectively; Extended Data Fig. 3)^11,31^. Given that human p97–UN requires longer or branched Ub chains for efficient unfolding compared with yeast Cdc48–UN^31,32^, we used the M1/K48-substrate for the following assays. The L53R and E84A mutations of UFD1 severely reduced p97–UN unfolding activity toward the M1/K48-substrate and abolished binding to K48-Ub ^init^ (Fig. 1c, e and Extended Data Fig. 4). These results confirm that Ub^prox^ binding is essential for p97–UN-mediated unfolding.

The Ub^init^ C-terminus (residues 68–73) forms an intermolecular β-sheet with UFD1 (residues 185–190), stabilized by a polar network (Fig. 1f). To validate the functional significance of these observed interactions, we assessed the unfolding activity of p97–UN harboring the UFD1 mutations K139A, E143A, E184A, or D186A (Fig. 1c). Except for E184A, all these mutations markedly reduced the unfolding efficiency (Fig. 1e). While E143A abolished binding to K48-Ub ^init^, the D186A mutant maintained interaction but failed to promote unfolding, suggesting that Asp186 participates in critical interactions with Ub^init^ before the initiation of unfolding. Additionally, a hydrophobic pocket of the Nc subdomain accommodates a C-terminal Leu cluster of Ub^init^ (Leu67, 69, 71, 73) that is buried in the native fold (Fig. 1g). While the exposure of these hydrophobic residues destabilizes the protein structure and promotes spontaneous refolding, the association of UFD1-UT3 with this leucine cluster stabilizes the unfolded conformation, thereby blocking refolding.

Our findings demonstrate that UFD1-UT3 engagement of both Ub^prox^ and Ub^init^ is indispensable for initiating unfolding. We propose a cooperative model where UFD1 and NPL4 synergistically anchor Ub^init^, with NPL4 and UFD1 capturing the N-terminal region (residues 1–48) and the C-terminal region (residues 66–76) of Ub^init^, respectively (Fig. 1h). This coordinated mechanism prevents refolding by shielding critical hydrophobic elements, maintaining Ub^init^ in an unfolded state primed for translocation through the p97 pore.

### UAS domain of UBXD7 and HUE motif of FAF1 and FAF2 Enhance p97–UN Activity

To identify additional p97 adaptors beyond the UN complex, we screened various UBX-domain-containing adaptors but found that none independently promoted p97 unfolding activity (Fig. 2a, b). However, further analysis revealed that FAF1, FAF2 lacking the membrane-anchor domain (FAF2^ΔMA^), and UBXD7 specifically enhance p97–UN unfolding activity (Fig. 2c). These adaptors partially rescued the activity impaired by UFD1 mutations (E84A, E143A, D186A), suggesting they can compensate for weakened UFD1–Ub interactions (Fig. 2d). Although these adaptors share a UBA–UAS-UBX architecture, AF3 predictions revealed structural divergence, with FAF1 and FAF2 featuring an extended ∼90-residue helix between the UAS and UBX domains, whereas UBXD7 contains a long flexible linker (Fig. 2e and Extended Data Fig. 5a, b).

**Figure 2.**
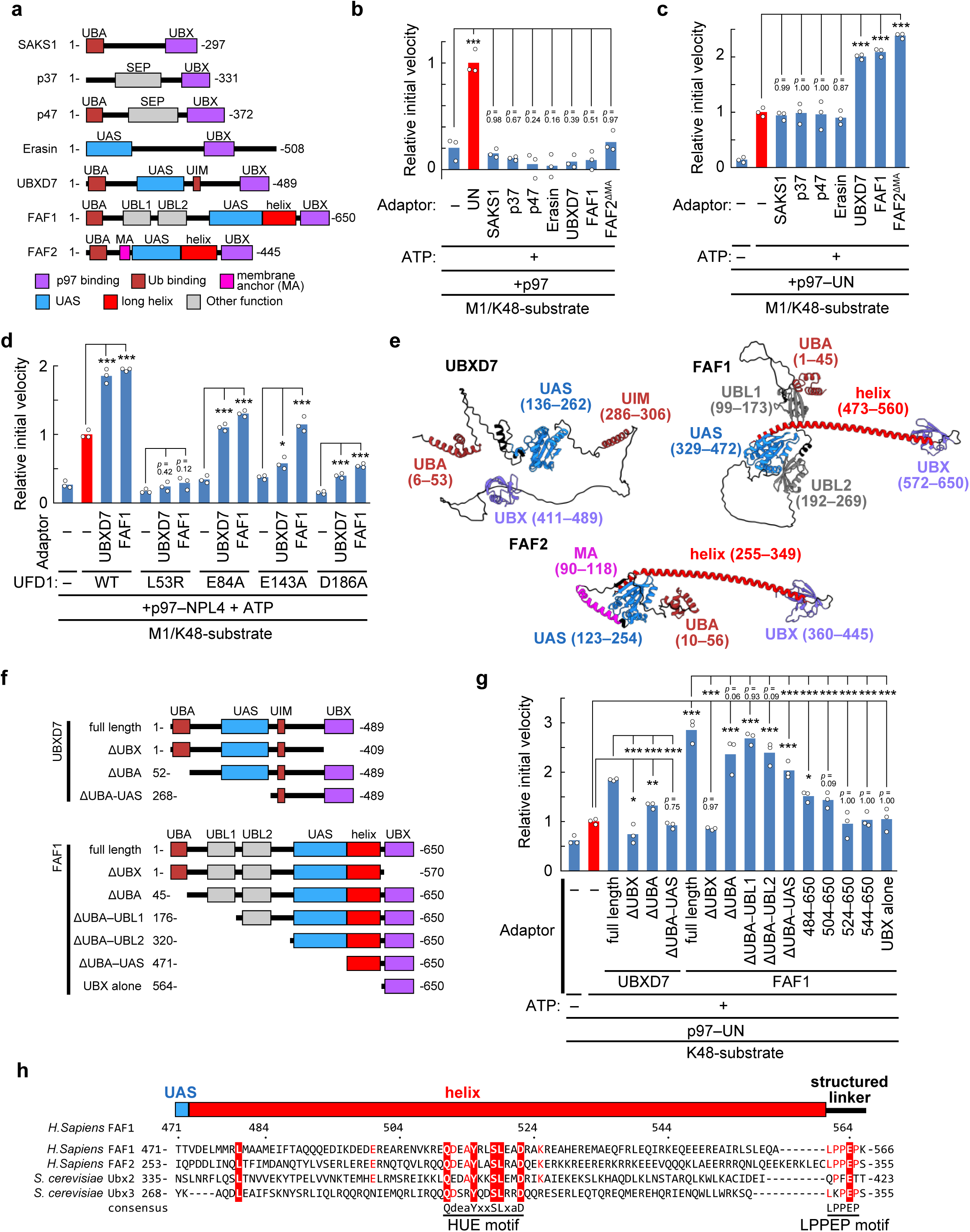
UBX-domain-containing adaptors FAF1, FAF2, and UBXD7 enhance p97–UN unfolding activity. **a,** Domain architectures of the UBX-domain-containing p97 adaptors used in this study. **b, c,** Impact of p97 adaptors on the unfolding activity of p97 (b), and p97–UN (c) toward the M1/K48-substrate. **d,** Impact of UBXD7 or FAF1 on the unfolding activity of p97–UN harboring various UFD1 mutants toward the M1/K48-substrate. **e,** Structural models of UBXD7, FAF1, and FAF2 predicted by AF3. Domains are colored according to the scheme in a. PAE plots for these predictions are provided in Extended Data Fig. 5a. **f,** Schematic diagrams of UBXD7 and FAF1 deletion mutants. **g,** Impact of the indicated deletion mutants of UBXD7 or FAF1 on the unfolding activity of p97–UN toward the K48-substrate. In **b–d**, and **g**, individual data points are shown as white dots, and bars represent the mean (n = 3 independent experiments). Relative initial velocities were normalized to the mean of the p97–UN complex in the presence of ATP (red bar). Statistical significance was determined using one-way ANOVA followed by Dunnett’s *post hoc* tests. \*\*\**p* < 0.001; \*\**p* < 0.01; \**p* < 0.05. Source data are provided in Extended Data Fig. 4. **h,** Sequence alignment of the extended helix connecting the UAS and UBX domains in FAF1, FAF2, and their yeast homologs, Ubx2 and Ubx3. Identical residues across all four sequences are highlighted in white text on a red background, while residues conserved in three sequences are shown in red. The HUE motif, defined by the highly conserved QdeaYxxSLxaD sequence (x represents any amino acid, lowercase letters indicate residues that are not conserved in the yeast), and the LPPEP motif are indicated below the alignment.

To identify the regions required for enhancement of the p97–UN activity, we analyzed deletion mutants of FAF1 and UBXD7 (Fig. 2f). Unfolding assays were performed using the K48-substrate, as the presence of FAF1 or UBXD7 stimulated a readily quantifiable unfolding response. Deletion analysis demonstrated that the UBX domain is indispensable for all three adaptors (Fig. 2g). However, their unfolding enhancer regions are distinct. Although UBA deletion attenuates UBXD7 activity, the UAS-UBX domains retain the ability to enhance p97–UN activity, whereas FAF1 requires the long helix between the UAS and UBX domains rather than the UBA and UAS domains. Fine-mapping of this FAF1 helix identified a core enhancer region (residues 504–523) containing a highly conserved QdeaYxxSLxaD sequence. We named this element the Helical Unfolding Enhancer (HUE) motif (Fig. 2h). The evolutionary conservation of the HUE motif from yeast to humans highlights its fundamental role in p97-mediated unfolding.

### Adaptor interactions with the UFD1-UT3 module stabilize the active unfolding assembly

We hypothesized that the p97–UN activity-enhancing regions of UBXD7, FAF1, and FAF2 function through protein-protein interactions. Accordingly, We used AF3 to predict the interactions of UBXD7, FAF1, and FAF2 with p97, NPL4, UFD1, and K48-chains. These factors constitute the essential components of the unfolding machinery. Because AF3 via the AlphaFold Server does not directly support K48-chains, we provided five separate Ub sequences as input. Based on the predicted models and PAE scores, the UBX domains of UBXD7, FAF1, and FAF2, which are essential for enhancing p97–UN activity, were shown to bind to the p97-NTD in a manner consistent with their known crystal structures^33,34^ (Extended Data Fig. 6a). In contrast, the UBXD7-UAS, FAF1-HUE, and FAF2-HUE specifically interact with UFD1-UT3. Furthermore, for UBXD7, the intrinsically disordered region between the UBA and UAS domains (residues 78–90, hereafter referred to as pre-UAS) also exhibited reduced expected positional error values relative to UFD1-UT3, suggesting its involvement in the interaction. Although FAF2-HUE was previously proposed as a Ub-binding motif^7^, the AF3 PAE plot exhibited expected positional errors exceeding 20 Å. Therefore, this interaction could not be predicted, at least in our attempts. Next, to determine whether UBXD7, FAF1, and FAF2 associate with UFD1-UT3 while it is engaged with K48-Ub ^init^, we performed AF3-based structural modeling of complexes comprising each accessory adaptor, UFD1, and K48-Ub ^init^ (consisting of a Ub and a C-terminal Ub^init^ fragment). The resulting PAE plots demonstrated that the inclusion of K48-Ub ^init^ further increased confidence in the interactions between the accessory adaptors and UFD1-UT3 (Fig. 3a-c and Extended Data Fig. 6b). These findings suggest that multivalent interactions among the adaptors, UFD1-UT3, and the K48 chain stabilize the active unfolding assembly.

**Figure 3.**
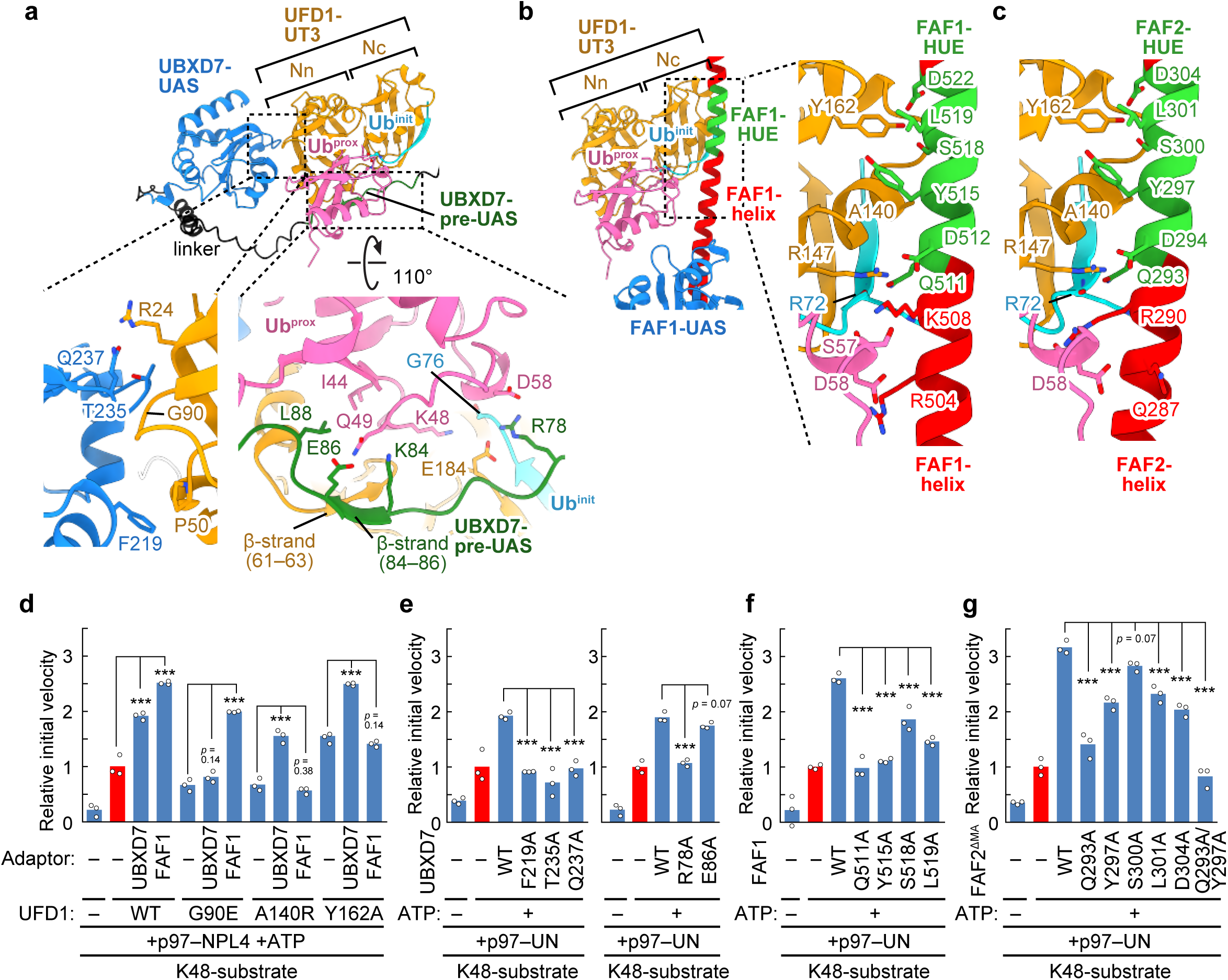
Structural models and functional validation of the interactions between UFD1-UT3–K48-Ub ^init^ and UBXD7, FAF1, or FAF2. **a–c,** AF3-predicted structures of UFD1–K48-Ub_2_^init^ in complex with UBXD7 (a), FAF1 (b), and FAF2 (c). UFD1-UT3 are oriented in the same direction to facilitate structural comparison. Regions not involved in the interactions are omitted for clarity. Because the overall structure of the FAF2 complex was nearly identical to that of the FAF1 complex, only close-up views are presented for the FAF2 complex. Color coding for UFD1–K48-Ub ^init^ follows the scheme in Fig. 1a, and the colors for UBXD7, FAF1, and FAF2 follow the scheme in Fig. 2a, with the pre-UAS region and HUE motifs highlighted in green. PAE plots for these predictions are provided in Extended Data Fig. 6b. **d,** Impact of UBXD7 or FAF1 on the unfolding activity of p97–UN harboring various UFD1 mutants toward the K48-substrate. **e–g**, Unfolding activity of p97–UN toward the K48-substrate in the presence of UBXD7 (e), FAF1 (f), or FAF2 (g) mutants. In **d–g**, individual data points are shown as white dots, and bars represent the mean (n = 3 independent experiments). Relative initial velocities were normalized to the mean of the p97–UN complex in the presence of ATP (red bar). Statistical significance was determined using one-way ANOVA followed by Dunnett’s *post hoc* tests. \*\*\**p* < 0.001. Source data are provided in Extended Data Fig. 4.

### Mechanism of p97-UN enhancement by the UBXD7 UAS and pre-UAS domains

The AF3-predicted structure of the UBXD7–UFD1–K48-Ub ^init^ complex reveals that the UAS domain and a newly identified pre-UAS region interact with the UFD1-UT3 Nn subdomain and Ub^prox^ (Fig. 3a). Hydrophobic and polar contacts involving UBXD7 residues Phe219, Thr235, and Gln237 and UFD1–UT3 residues Arg24, Pro50 and Gly90 stabilize the UBXD7-UAS–UFD1 interface. The G90E mutation in UFD1, or the F219A, T235A, Q237A substitutions in UBXD7, completely abolished the enhancement of p97–UN activity for the K48-substrate by UBXD7 (Fig. 3d, e). While the pre-UAS of UBXD7 is an intrinsically disordered region in the absence of UFD1-UT3, residues 84-86 in pre-UAS form a β-sheet with residues 61-63 of UFD1-UT3 (Fig. 3a). Furthermore, Arg78, Lys84, and Glu86 within the UBXD7 pre-UAS form polar interactions with UFD1 and Ub^prox^. In addition, Leu88 of UBXD7 is positioned near the canonical Ile44 hydrophobic patch of Ub^prox^. While the E86A mutation in UBXD7 had a negligible impact on activity, the R78A substitution abolished the UBXD7-mediated enhancement of p97-UN activity (Fig. 3e). Because the binding affinity between UFD1-UT3 and K48-chains is nearly undetectable before the initiation of unfolding (Fig. 1c), these findings suggest that UBXD7 promotes the initiation of the unfolding process by stabilizing this transient association between UFD1-UT3 and the K48-chain.

Unlike UBXD7, the UAS domains of FAF1, FAF2, and Erasin fail to enhance p97–UN activity (Fig. 2). Sequence comparison reveals that these adaptors lack the essential residues for UFD1 binding (Extended Data Fig. 7a). We also used AF3 to analyze the binding of the UAS domains of UBXD7, FAF1, FAF2, and Erasin to UFD1-UT3. Their PAE plots exhibited expected positional errors exceeding 15 Å, indicating an inability to interact with UFD1-UT3 (Extended Data Fig. 7b). These findings demonstrate that the UAS domains of other adaptors lack the necessary structural elements to associate with UFD1. Conversely, Ubx5, the yeast homolog of UBXD7, retains Arg78 in pre-UAS and Phe219 and Thr235 in the UAS domain. In Ubx5, Gln237 is replaced by glutamate, a residue that retains the ability to form equivalent interactions. This conservation highlights the structural uniqueness and evolutionary significance of the UBXD7 pre-UAS and UAS domains.

### FAF1- and FAF2-helices Enhance p97–UN Activity through their binding to UFD1-UT3 and Ub moieties

The AF3-predicted structure of the FAF1–UFD1–K48-Ub ^init^ complex reveals that the FAF1-HUE motif interacts with Ala140, Arg147, and Tyr162 residues of UFD1-UT3 (Fig. 3b). Furthermore, Gln511 in FAF1-HUE is positioned to hydrogen-bond with the Ub^init^ main-chain CO group. Mutational analyses confirmed the importance of these interactions. The A140R and Y162A mutations in UFD1 abolished the FAF1-mediated enhancement of p97–UN activity (Fig. 3d). Likewise, S518A and L519A mutations in FAF1-HUE reduced activity while Q511A and Y515A mutations completely abolished unfolding enhancement (Fig. 3f). These residues in FAF1-HUE are conserved in FAF2-HUE, in which Q293A, Y297A, L301A, and D304A mutations significantly impaired its function to enhance p97–UN activity, while the Q293A/Y297A double mutant resulting in a complete loss of enhancement (Fig. 3g). Additionally, despite the lack of sequence conservation between the N-terminal extensions of the HUE motifs of FAF1 and FAF2, this region is predicted to engage in polar interactions with Ub^prox^, thereby facilitating complex stabilization (Fig. 3b, c).

While HUE motifs have been shown to bind Ub^init^, we performed additional predictions to determine whether the HUE motif also engages Ub in its folded state before unfolding. AF3 structural prediction of a complex comprising the FAF1-helix, UFD1-UT3, and two full-length Ub sequences revealed that Gln511 of FAF1 interacts with folded Ub^init^, similar to the arrangement observed in the FAF1–UFD1–K48-Ub ^init^ model (Extended Data Fig. S7c, d). Notably, the addition of the FAF1-helix stabilized the ensemble by suppressing structural variability in Ub^dist^ and marginally improving the PAE scores for Ub^init^ (Extended Data Fig. 1a, 7c, e). This increased structural consistency likely results from FAF1-HUE triggering a transition of the Ub^init^ C-terminus from an intramolecular β-sheet to an intermolecular β-sheet with UFD1-UT3, as also observed in the structure of UFD1-UT3-K48Ub2init (Fig. 1a). This facilitated assembly of Ubinit with UFD1-UT3 represents a key mechanism for FAF1-mediated enhancement of unfolding.

To investigate whether the interaction between the HUE motif and UFD1-UT3 is required for cellular function, we analyzed the inhibitory effect of FAF2 on pexophagy^27^ (Fig. 4a, b). While FAF2 deficiency promotes pexophagy, reintroduction of FAF2 WT suppresses this process. However, pexophagy remained suppressed to a level comparable to the WT by the Q293A/Y297A double mutant, despite its near-complete loss of p97–UN unfolding enhancement activity (Fig. 3g). In contrast, the FAF2^Δhelix^ mutant (Δ275–350) was less effective in suppressing pexophagy. Furthermore, recent findings show that FAF2 triple mutations (L289N, Y297H, L301N) partially impair inhibition of pexophagy^7^. This variant likely exerts a stronger effect than our double mutant because it incorporates L289N, which targets the Ub^prox^-interacting region.

**Figure 4.**
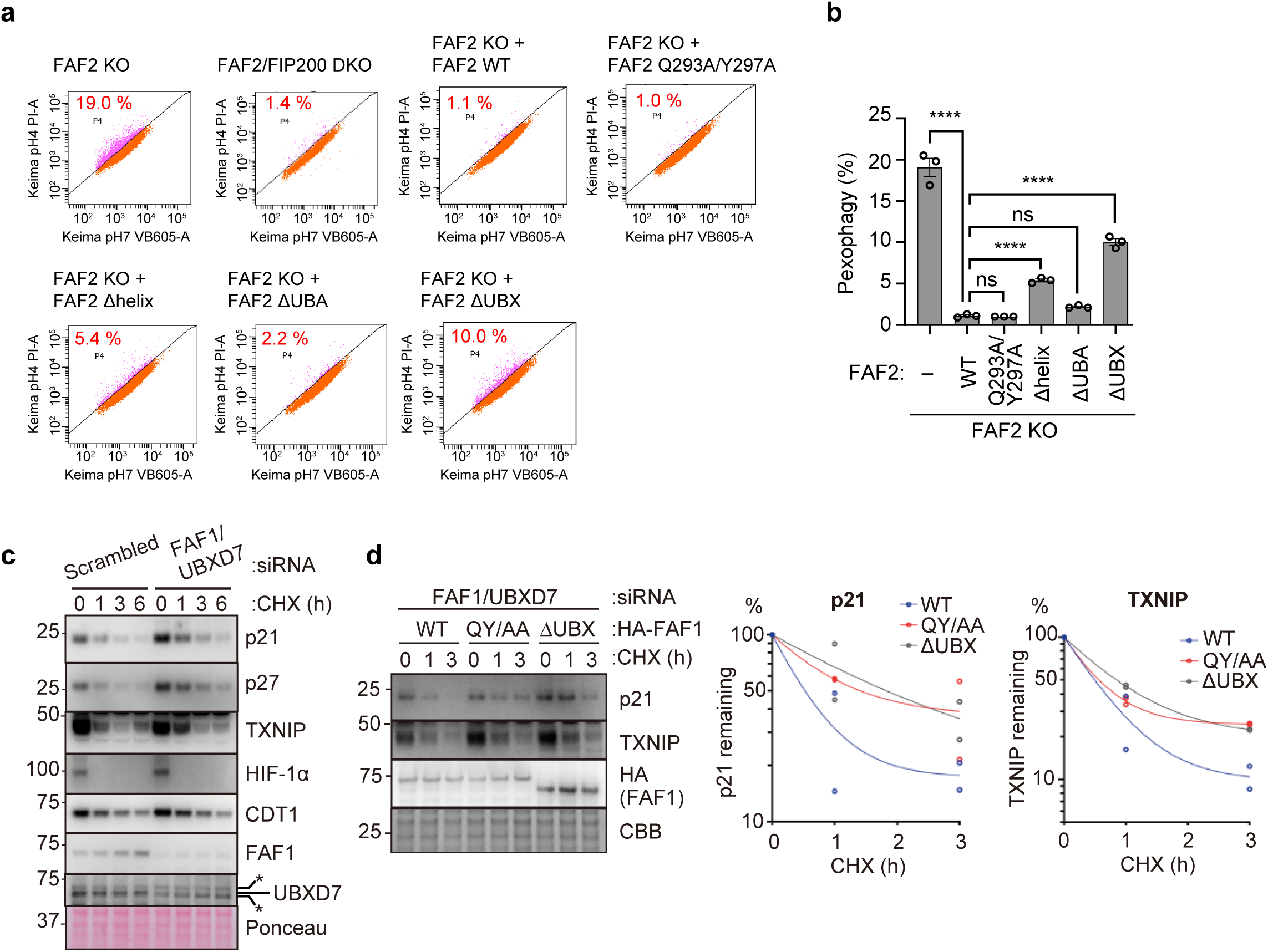
Functional roles of the FAF1 and FAF2 HUE motifs in cellular protein degradation and pexophagy. **a,** FACS-based analysis of the pexophagy flux. Representative FACS data (mKeima-SKL 561/488 ratio) for WT, *FAF2* KO, or *FAF2* KO stably expressing the indicated 3HA-FAF2 mutants, with the percentage of pexophagy-positive cells indicated. **b,** Quantitative analysis of the pexophagy flux for cells from a. Data represent the mean percentage of pexophagy-positive cells from three independent experiments. Error bars represent mean ±SEM. Statistical significance was determined using one-way ANOVA with Tukey’s multiple comparisons test. \*\*\*\**p* < 0.0001; ns. not significant. **c**, Cycloheximide (CHX) chase assay in HEK293T cells transfected with the indicated siRNAs for 72 h and treated with CHX for 0–6 h. Proteins were detected by western blotting using the indicated antibodies. **d,** CHX chase assay in HEK293T cells depleted of FAF1 and UBXD7 by siRNA (72 h) and rescued with HA-tagged WT, QY/AA (Q511A/Y515A), or ΔUBX mutant FAF1 (48 h). Cells were treated with CHX for 0–3 h. Proteins were analyzed by western blotting using the indicated antibodies. Graphs on the right show quantification of protein levels normalized to the signal at 0 h, defined as 100% (p21 or TXNIP; n = 2 biological replicates).

Notably, the persistence of pexophagy suppression in the FAF2^ΔUBX^ mutant implies that FAF2 operates through an additional p97-independent pathway, thereby explaining why the FAF2^Δhelix^ mutant also retains its inhibitory effect.

We subsequently performed cycloheximide (CHX) chase assays using FAF1. siRNA-mediated knockdown of FAF1 and UBXD7 reduced endogenous FAF1 and UBXD7 levels and delayed the degradation of p97 clients, such as p21 and TXNIP, following CHX treatment (Fig. 4c). In rescue experiments, reintroduction of HA-FAF1 WT restored the degradation of p21 and TXNIP. In contrast, the HA-FAF1 Q511A/Y515A (QY/AA) mutant exhibited delayed degradation at a level comparable to that of FAF1^ΔUBX^ (Fig. 4d). These findings indicate that the FAF1-HUE–UFD1-UT3 interaction serves as a central hub for promoting protein degradation within the cellular environment.

### Non-cooperative Activation of p97–UN by FAF1 and UBXD7

Unfolding assays revealed that p97–UN harboring the UFD1 G90E was stimulated exclusively by FAF1, while p97–UN harboring the UFD1 A140R or Y162A was enhanced specifically by UBXD7 (Fig. 3d). These results confirm that FAF1-HUE and UBXD7-UAS occupy distinct binding sites on UFD1-UT3. Although the AF3 model of the FAF1–UBXD7–UFD1–K48-Ub_2_^init^ complex indicates that simultaneous binding is structurally possible, biochemical assays showed no synergistic enhancement of p97-UN activity by the combination of FAF1 and UBXD7 (Extended Data Fig. 8a-c). This lack of synergy likely arises from both FAF1-UBX and UBXD7-UBX targeting the same p97-NTD. To assess the occupancy of multiple p97–NTDs by the UBX domain, we analyzed the binding stoichiometry with p97–UN using size-exclusion chromatography (SEC). Consistent with previous reports^34^, our analysis revealed a 6:1:1 stoichiometry for both the p97–UN–FAF1 and p97–UN–UBXD7 complexes (Extended Data Fig. 8d). This result suggests that these UBX domains selectively target a specific NTD of the p97 hexamer. Due to their structural similarity, both UBX domains likely target the same specific p97-NTD, causing the adaptors to compete for recruitment. This mutual exclusivity prevents cooperative activation and allows only one adaptor to engage the complex at a time.

### Structural principles of FAF1-HUE positioning and coordinated p97-UN activation

We employed AF3 to predict the architecture of the p97–UN–Ub^init^–FAF1 complex. Due to the 5,000-residue input limit of the AlphaFold server^28^, we used six chains of p97 consisting solely of the NTD and D1 domains (residues 1-463, designated as p97^ND1^), along with single chains of UFD1, NPL4, FAF1, and a C-terminal Ub^init^ fragment (residues 67–76). We completed the model by superimposing residues 13–50 of Ub^init^ from the Cdc48-UN cryo-EM structure (PDB ID 8DAV)^19^. In this model, FAF1-UBX binds the specific p97-NTD already engaged by the UFD1-SHP1 motif (Fig. 5a, b, Extended Data Fig. 9a). The linker connecting the helix and the UBX domain in FAF1 is stabilized by interactions between its internal LPPEP motif and the UBX domain. These residues are uniquely conserved in FAF1 and FAF2 (Fig. 5c). This structural arrangement ensures that the helix containing the HUE motif is positioned above the p97 hexameric ring. This model matches the previously reported cryo-EM density of the 6:3 complex of p97–FAF1 (EMD-2319)^35^, validating our prediction (Extended Data Fig. 9b). Furthermore, FAF1-HUE was found to bind UFD1-UT3, thereby positioning the C-terminal fragment of Ub^init^ close to its N-terminal segment in the NPL4-tower (Fig. 5a).

**Figure 5.**
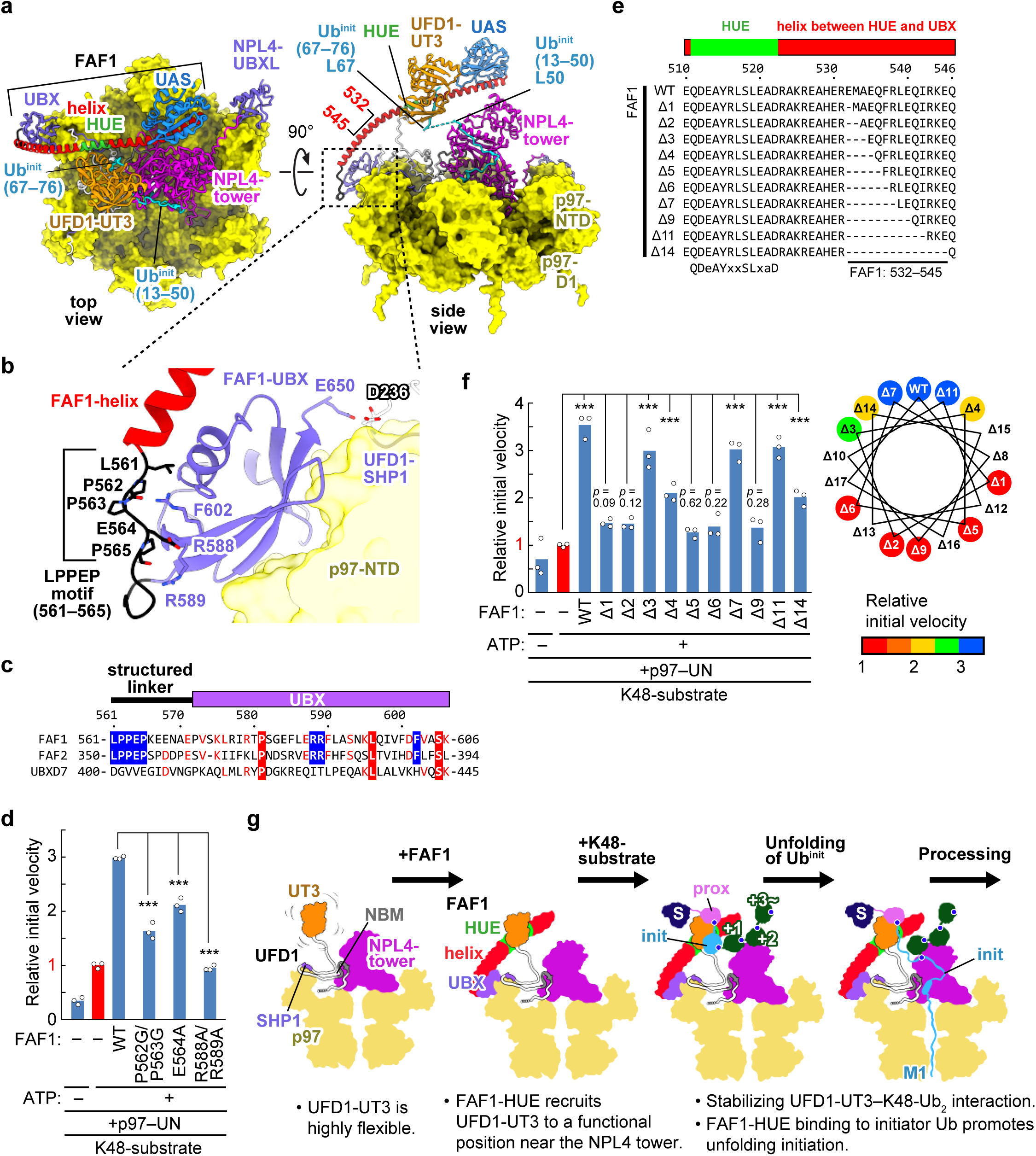
Mechanism of p97–UN activity enhancement by FAF1. **a,** Structural model of the p97–UN–Ub^init^–FAF1 complex generated by integrating AF3 predictions and cryo-EM data (PDB ID: 8DAV)^19^. The p97-D2 ring is omitted due to the residue input limit of the AlphaFold server. The flexible N-terminal region of FAF1 preceding the UAS domain and the C-terminal NZF domain of NPL4 are removed for clarity. p97 is shown as a surface model, while other components are presented as cartoon models. Color coding for UFD1–UT3 and Ub^init^ follows Fig. 1a; other regions follow Fig. 2a, with the FAF1-HUE motif in green, p97 in yellow, and the NPL4-tower in magenta. The PAE plot for this prediction is in Extended Data Fig. 9a. **b,** Close-up view of the interactions involving FAF1-UBX, the p97-NTD bound to UFD1-SHP1, and the LPPEP motif. **c,** Sequence alignment of FAF1, FAF2, and UBXD7 from the LPPEP motif to the N-terminal region of the UBX domain. Identical residues are in white on a red background, and those conserved in two sequences are in red. The LPPEP motif and its predicted interacting residues are highlighted in white text on a blue background. **d,** Unfolding activity of p97–UN in the presence of FAF1 variants targeting the intramolecular LPPEP–UBX interaction interface. **e,** Amino acid sequences of FAF1 mutants with shortened helices between the HUE motif and the UBX domain. **f,** Unfolding activity of p97–UN in the presence of helix-shortening FAF1 mutants. The helical wheel illustrates the shift in the UFD1-UT3 binding site for each mutant, with colors indicating the resulting enhancement of p97–UN activity. In **d** and **f**, individual data points are shown as white dots, and bars represent the mean (n = 3 independent experiments). Relative initial velocities were normalized to the mean of the p97–UN complex in the presence of ATP (red bar). Statistical significance was determined using one-way ANOVA followed by Dunnett’s *post hoc* tests. \*\*\**p* < 0.001. Source data are provided in Extended Data Fig. 4. **g,** Proposed model for FAF1-mediated enhancement of p97–UN activity. Schematic representations follow the style in Fig 1h. The FAF1 helix, HUE motif, and UBX domain are colored red, green, and purple, respectively.

To confirm that FAF1-HUE positioning is critical for p97-UN activation, we disrupted the LPPEP–UBX interface. While P562G/P563G and E564A mutations reduced activity, the R588A/R589A mutation completely abolished the FAF1-mediated enhancement of p97–UN (Fig. 5b, d). Disruption of the LPPEP–UBX interface likely causes the HUE motif to shift from its optimal position, demonstrating that precise HUE positioning is essential for p97-UN activation.

Next, we generated FAF1 helix deletion mutants to evaluate how HUE orientation impacts p97-UN activity (Fig. 5e, f). Each single-residue deletion rotates the UFD1-UT3 binding site of HUE by 100 degrees and shortens the helix by 1.5 Å. Orientations opposite to WT (Δ1, 2, 5, 6, or 9) abolished activity, while those restoring WT-like orientations (Δ3, 4, 7, 11, or 14) preserved enhancement. Notably, FAF2 maintains this precise geometry through a seven-residue insertion that yields a helical orientation equivalent to an 11-residue deletion (Fig. 2h, 5f). While p97–UN activity is highly sensitive to orientation shifts of FAF1-HUE, it remains robust to spatial shifts up to 16.5 Å (Δ11). In contrast, a 21 Å reduction (Δ14) of the FAF1-helix impaired the ability to enhance p97–UN activity, likely because the excessive distance between UFD1-UT3 and the NPL4-tower hinders their coordinated action.

To maintain precise HUE orientation, FAF1-UBX must selectively target the p97-NTD already engaged by UFD1-SHP1 (Fig. 5a, b). Although previous reports indicated that FAF1-UBX can associate with up to three p97-NTDs in the absence of UN, our prediction that FAF1-UBX binds to a specific p97–NTD conforms to SEC analysis, which demonstrated that the p97–UN–FAF1 complex assembles with a 6:1:1 stoichiometry (Extended Data Fig. 8d)^34,35^. While AF3 predicted FAF1 Glu650 to be near UFD1, the E650A mutation did not affect activity (Extended Data Fig. 9c). This suggests that direct FAF1–UFD1 interaction does not determine the binding site. Instead, since the UFD1-bound NTD exclusively adopts an “Up” conformation^36,37^, this structural state likely serves as the recruitment signal that ensures FAF1-UBX binds the correctly positioned p97-NTD.

Our findings establish a mechanistic framework for how FAF1 and FAF2 enhance p97–UN activity. Although UFD1-UT3 is naturally flexible^13–16,38^, FAF1-HUE serves as a scaffold to recruit it near the NPL4-tower (Fig. 5g). This spatial arrangement enables UFD1-UT3 and NPL4 to coordinate Ub^init^ unfolding. The FAF1-helix further stabilizes the complex via simultaneous interactions with UFD1-UT3 and the K48-chain (Fig. 3b). Crucially, FAF1-HUE may accelerate the remodeling of the Ub^init^ intramolecular β-sheet into an intermolecular one with UFD1-UT3 (Extended Data Fig. 7e). We propose a dual mechanism model. FAF1 and FAF2 drive unfolding by strategically positioning UFD1-UT3 and directly promoting this rate-limiting β-sheet exchange.

### Divergent structural strategies for p97-UN activation by FAF1 and UBXD7

Following the strategy employed for FAF1, we performed AF3-based structural modeling of the p97–UN–Ub^init^–UBXD7 complex (Fig. 6a and Extended Data Fig. 9d). Similar to FAF1, UBXD7-UBX is predicted to bind the p97-NTD already engaged by UFD1-SHP1. However, because the D487A/N489A double mutation in the UBXD7-UBX near UFD1-SHP1 retains wild-type activity, UBXD7-UBX likely recognizes the “Up” conformation of the p97-NTD for its recruitment. Crucially, the AF3 prediction that both UBXD7-UBX and FAF1-UBX bind to the same p97-NTD is consistent with the experimental observation that FAF1 and UBXD7 do not synergistically enhance p97–UN activity (Extended Data Fig. 8c).

**Figure 6.**
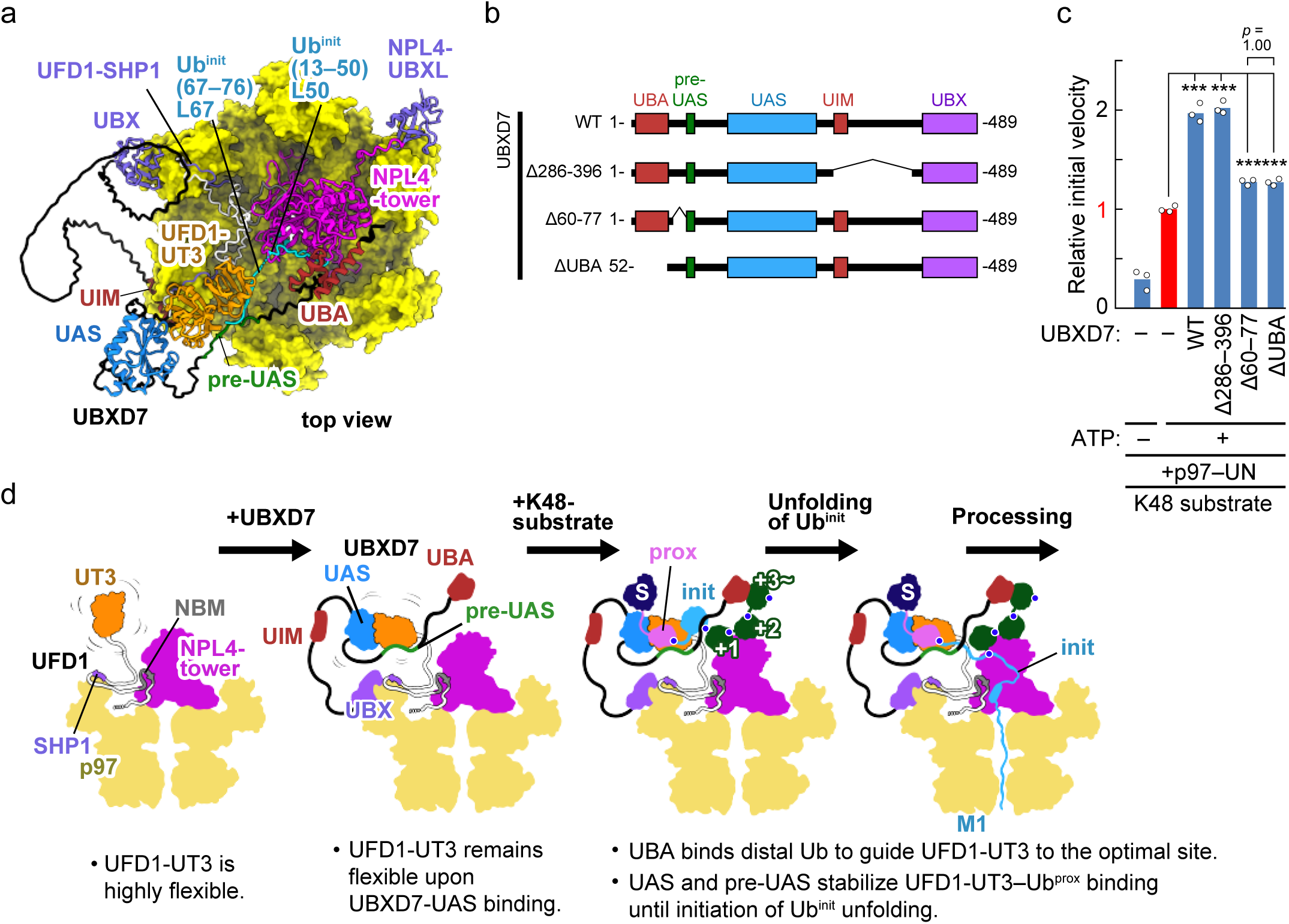
Mechanisms of p97–UN activity enhancement by UBXD7. **a,** Structural model of the p97–UN–Ub^init^–UBXD7 complex generated by integrating AF3 predictions and cryo-EM data (PDB ID: 8DAV)^19^. Except for UBXD7, all components are shown as in Fig. 5a. UBXD7 is colored according to the scheme in Fig. 2a, with the pre-UAS region highlighted in green. The PAE plot for this prediction is provided in Extended Data Fig. 9d. **b,** Schematic diagrams of UBXD7 deletion mutants. **c,** Unfolding activity of p97–UN in the presence of UBXD7 deletion mutants. Individual data points are shown as white dots, and bars represent the mean (n = 3 independent experiments). Relative initial velocities were normalized to the mean of the p97–UN complex in the presence of ATP (red bar). Statistical significance was determined using one-way ANOVA followed by Dunnett’s *post hoc* tests. \*\*\**p* < 0.001. Source data are provided in Extended Data Fig. 4. **d,** Proposed model for UBXD7-mediated enhancement of p97–UN activity. Schematic representations follow the style of Fig. 1h. UBXD7 is colored as in Fig. 1a, with the pre-UAS highlighted in green.

Despite their shared UBX binding mode, FAF1 and UBXD7 exhibit significant divergence outside their respective UBX domains. UBXD7 lacks the rigid helix present in FAF1, resulting in greater flexibility of the UAS domain. This structural difference suggests that UBXD7 functions through a mechanism distinct from the FAF1-mediated positioning of UFD1-UT3. Notably, a UBXD7 variant lacking the flexible UIM-containing linker (residues 286–396) retained WT-level of p97–UN activity enhancement (Fig. 6b, c). Thus, unlike in FAF1, the region between the UAS and UBX domains of UBXD7 is not essential for enhancing activity.

UBXD7 and FAF1 exhibit divergent functional requirements for their respective UBA domains. While the ΔUBA mutation in FAF1 exerts a negligible effect, it attenuates UBXD7-mediated enhancement of p97–UN activity (Fig. 2g). As the initiation of unfolding necessitates the engagement of Ub^prox^ and Ub^init^ by UFD1-UT3 (Fig. 1d, g), the UBA domain is sterically hindered from binding these specific subunits and likely associates with Ub^dist^ moieties instead. Notably, deletion of the linker between UBA and pre-UAS (residues 60–77) reduced activity to levels observed in the ΔUBA mutant (Fig. 6b, c). These findings indicate that UBA-mediated tethering to Ub^dist^, coordinated by the linker, properly positions the UAS domain to ensure efficient substrate processing (Fig. 6d). This UBA-directed recruitment of the UAS domain toward Ub^prox^ likely facilitates K48-chain capture by the pre-UAS and UAS domains, thereby stabilizing the complex until the onset of Ub^init^ unfolding (Fig. 3a).

## Discussion

Our 1.31 Å crystal structure of the UFD1-UT3-K48-Ub ^init^ complex identifies the UT3 domain as the pivotal module for initiating p97-mediated unfolding (Fig. 1a, h). Although UFD1 exhibits modest affinity for native K48-chains, it transitions to exceptionally high affinity upon Ub^init^ unfolding (Fig. 1c), ensuring stable engagement throughout the reaction. AF3-based prediction of the unfolded Ub^init^ state facilitated our crystallographic analysis, revealing that the Nc subdomain groove specifically recognizes the C-terminal segment of Ub^init^ (Fig. 1f, g). This association stabilizes the unfolded conformation and likely represents the rate-limiting step, as sequestering typically buried hydrophobic residues enables efficient initiation.

Specialized adaptors FAF1, FAF2, and UBXD7 enhance p97 activity via divergent structural strategies that act through the UFD1-UT3 module. FAF1 and FAF2 adopt a rigid helix to scaffold UFD1-UT3 into a functional orientation near the NPL4-tower (Fig. 5g). Conversely, UBXD7 employs its UAS and pre-UAS domains to stabilize transient interactions between UFD1-UT3 and Ub^prox^ (Fig. 3a, 6d). Cellular assays confirm that the FAF1-HUE-mediated interaction is essential for protein homeostasis and efficient client degradation (Fig. 4d). This result illustrates how such regulatory modes accelerate substrate processing.

The discovery of the Nc subdomain groove provides a strategic advantage for therapeutic development (Fig. 1f, g). While p97 is a high-priority target for the treatment of cancer and other diseases^39^, the clinical utility of ATPase inhibitors is limited by low selectivity. A significant benefit of targeting UFD1 and NPL4 is that it enables specific inhibition of the UPS, thereby avoiding interference with the diverse non-UPS functions of p97. Although various small-molecule inhibitors and peptides targeting NPL4 have been developed^40,41^, UFD1-specific inhibitors remain unavailable, primarily because the molecular mechanisms underlying UFD1 function have been largely uncharacterized. The Ub^init^-binding groove identified in this work is essential for p97–UN activity and possesses a pocket architecture suitable for small-molecule engagement. Because this site is unique to the p97–UN complex, it represents a promising target for developing next-generation, UPS-specific p97 inhibitors. Such agents could offer a more precise approach to modulating protein degradation in human disease, providing a therapeutic strategy that moves beyond the limitations of current p97-directed therapies.

## Author contributions

Y.S. conceived the study and wrote the manuscript. Y.S. performed structural predictions. S.M., N.Y., and Y.S. performed biochemical and biophysical characterization. A.H. and Y.S. performed crystal structure determination. Y.G. and G.H. executed the chemical synthesis of Ub variants and performed BLI assays. Y.A. and F.O. performed CHX-chase assays. F.K. and N.M. conducted pexophagy assays. All authors analyzed the data and discussed the results.

## Notes

The authors declare no competing financial interest.

## Acknowledgments

We thank the beamline staff of the biological crystallography beamline of BL32XU of SPring-8 (Hyogo, Japan) for the technical help during data collection. We thank the BINDS staff for their support. This research was supported by Research Support Project for Life Science and Drug Discovery (Basis for Supporting Innovative Drug Discovery and Life Science Research (BINDS)) from AMED under Grant Number JP25ama121001 (support number 6703). This work was supported by JSPS/MEXT KAKENHI (grant numbers JP21H00283, JP21H02418, JP24H01899, and JP25H00977 to Y.S.; JP21K06161 to F.K.; JPMXP1323015483 to N.M.; JP23H04922 and JP25H00977 to F.O; JP24H01893 and 25H01300 to G.H.), MEXT Leading Initiative for Excellent Young Researchers (to Y.S.), Tottori University Research Support Project for the Next Generation (to Y.S.), TMDU priority research areas grant (to F.K.), MRI research grant FY2023 (to F.K.), the Naito Foundation (to F.K.), Female Faculty Promotion Program (FFPP) of Science Tokyo (to F.K.), AMED-CREST (grant number JP21gm1410007 to F.O.; JP20gm1410004 to N.M.), and AMED-BINDS (grant number JP22ama121009 to G.H.). During the preparation of this work, the authors used Gemini (Google) for English language editing and grammar correction. After using this tool, the authors reviewed and edited the content as needed and take full responsibility for the content of the publication.

## Data availability

The coordinates and structure factors of UFD1-UT3–K48-Ub ^init^ have been deposited in the Protein Data Bank under the accession code 24GI.

## METHODS

### Structural prediction

Structural prediction was performed using the AlphaFold Server powered by AlphaFold3 (https://alphafoldserver.com)^28^. The resulting structure models and Predicted Aligned Error (PAE) maps were visualized using ChimeraX^42^.

### Expression system and construction

The pET28a-SUMO plasmid was constructed by inserting the *Saccharomyces cerevisiae* SUMO sequence into the *NdeI* site of the pET28a plasmid, enabling the addition of an N-terminal His_6_-SUMO tag. The pET28a-SUMO plasmids containing *Escherichia coli* (*E. coli*) codon-optimized genes of human p97 and NPL4, as well as the pGEX-6P1 plasmid containing the codon-optimized gene of human UFD1, were prepared in a previous study^41^. The pET28a-SUMO plasmid containing C-terminal His_8_-tagged Ub_4_-mEos3.2 was also constructed previously, and the plasmid containing C-terminal His_8_-tagged Ub-mEos3.2 was generated by PCR. Genes encoding SAKS1, p37, p47, Erasin, UBXD7, FAF1, and FAF2 were PCR-amplified from Human Brain, Whole QUICK-Clone cDNA (Clontech). Since FAF2 is a membrane-anchored (MA) protein, we removed the MA region (residues 90–118) to create FAF2^ΔMA^ for soluble expression. The amplified genes were inserted into the pET28a-SUMO plasmid using the HiFi DNA Assembly Cloning Kit (New England Biolabs). Truncation and Site-directed mutations were introduced by PCR-based mutagenesis.

### Protein expression and purification

*E. coli* strain Rosetta 2 (DE3) cells (Novagen) transformed with various expression vectors were cultured in LB medium containing 50 mg L^-1^ ampicillin or 25 mg L^-1^ kanamycin at 37 °C. When the optical density at 600 nm reached approximately 0.5, isopropyl-β-D-thiogalactopyranoside (IPTG) was added to a final concentration of 0.1 mM to induce protein expression, and the cultures were incubated for an additional ∼18 h at 15 °C. Cells overexpressing N-terminal His_6_-SUMO-tagged proteins were harvested by centrifugation at 7000 × *g* for 10 min and disrupted by sonication in 50 mM Tris-HCl buffer (pH 8.0) containing 150 mM NaCl and 0.5% Triton X-100. The lysates were purified using a nickel-nitrilotriacetic acid (Ni-NTA) column (Qiagen). To remove the His_6_-SUMO tag, the eluted proteins were treated with Ulp1 protease for ∼15 h at 4 °C. The samples were further purified using a HiTrap Q HP anion exchange column (Cytiva). p97 was further purified by size-exclusion chromatography (SEC) on a Superose 6 column (Cytiva). Cells overexpressing N-terminal GST-tagged UFD1-UT3 (residues 17-196) were harvested by centrifugation at 7000 × *g* for 10 min and disrupted by sonication in PBS containing 1 mM DTT and 0.5% Triton X-100. The lysates were purified using a Glutathione Sepharose FF column (Cytiva). To remove the GST tag, the eluted protein was treated with HRV3C protease for ∼15 h at 4 °C. The samples were further purified using a Resource Q anion-exchange column (Cytiva) and SEC on a HiLoad Superdex 75 column (Cytiva).

To prepare the UFD1–NPL4 1:1 complex, *E. coli* strain Rosetta 2 (DE3) cells co-transformed with pGEX-6P1-UFD1 and pET28a-SUMO-NPL4 were cultured in LB medium containing 50 mg L^-1^ ampicillin and 25 mg L^-1^ kanamycin at 37 °C. When the optical density at 600 nm reached approximately 0.5, isopropyl-β-D-thiogalactopyranoside (IPTG) was added to a final concentration of 0.1 mM to induce protein expression, and the cultures were incubated for an additional ∼18 h at 15 °C. Cells overexpressing GST-UFD1 and His_6_-SUMO-NPL4 were harvested by centrifugation at 7000 × *g* for 10 min and disrupted by sonication in 50 mM Tris-HCl buffer (pH 8.0) containing 150 mM NaCl and 0.5% Triton X-100. The lysates were purified using a Glutathione Sepharose FF column. To remove the GST tag, the eluted GST-UFD1–His₆-SUMO-NPL4 complex was treated with HRV 3C protease for ∼15 h at 4 °C. The resulting UFD1–His₆-SUMO-NPL4 was purified by Ni-NTA affinity chromatography. To remove the His₆-SUMO tag, the eluted protein was further cleaved with Ulp1 protease for ∼15 h at 4 °C. Finally, UFD1–NPL4 was purified by SEC on an ENrich SEC 70 column (Bio-Rad).

### Chemical synthesis of biotinylated K48-Ub_2_ ^init^, biotinylated Ub and K48-Ub_2_ ^init^

For the preparation of peptide segments, automated solid-phase peptide synthesis was performed by using Initiator+ Alstra (Biotage) or Liberty blue (CEM) via the standard Fmoc-SPPS protocol. Coupling conditions are as follows; Fmoc-protected amino acids (4 equiv), HBTU (3.9 equiv) and DIEA (8 equiv) in DMF for 5 min at 75 °C for Biotage, and Fmoc-protected amino acids (4 equiv), DIC (8 equiv) and OxymaPure (4 equiv) in DMF for 2 min at 90 °C for CEM. Fmoc removal was conducted with 20% piperidine/DMF for 5 min at 50 °C twice for Biotage, and for 1 min at 80 °C for CEM. For the synthesis of peptide thioester, 2-chlorotrityl chloride resin was used followed by coupling of Fmoc-NHNH_2_ before peptide elongation, while Rink-amide PEG resin and Fmoc-Gly-Wang-PEG resin were used for peptides amide and acid, respectively. For C-terminal biotin labeling, Fmoc-Lys(Biotin)-OH was coupled as the C-terminal amino acid. Alloc-Lys(Fmoc)-OH was introduced at the branching point (i.e. K48) and Alloc group was removed after the first chain elongation was completed. After completion of peptide elongation, peptide cleavage and global deprotection were performed using TFA/TIS/H₂O (95/2.5/2.5 for peptide hydrazide and acid) or TFA/TIS/ DMB (92.5/2.5/5 for peptide amide) for 2 h. After purification of crude peptides by preparative HPLC, each peptide was analyzed by analytical HPLC for purity check, and was identified by MALDI-TOF MS.

For peptide ligation, peptide hydrazide was dissolved in a buffer containing 6 M guanidine hydrochloride (Gn·HCl) and 0.2 M NaH₂PO₄ (pH 3.0). Acetylacetone (10 equiv.) was added and the pH was adjusted to 2.8–3.0. After completion of C-terminal activation, MPAA was added (final conc. 100 mM), and the pH was adjusted to 5.8–6.0. Upon completion of the thioesterification, the N-terminal Cys peptides (0.8–1.0 equiv) and TCEP (final conc. 50 mM) were directly mixed. The pH was adjusted to 6.0–6.5, and the reaction mixture was stirred at 37 °C. The reaction progress was monitored by analytical HPLC and MALDI-TOF MS. After purification of ligated products by preparative HPLC, the isolated products were analyzed by analytical HPLC for purity check, and identified by MALDI-TOF MS.

For desulfurization, ligated peptide was dissolved in a degassed solution of 6 M guanidine hydrochloride (Gn·HCl) and 0.2 M NaH₂PO₄ buffer (pH 6.9). Then, TCEP, glutathione and VA-044 were added to give final concentrations of 0.5 mM peptide, 300 mM TCEP, 80 mM glutathione, and 20 mM VA-044. The pH of the reaction mixture was adjusted to 6.7–7.0, and the mixture was stirred at 37 °C for 12 hours under an argon atmosphere. After purification of ligated products by preparative HPLC, the isolated products were analyzed by analytical HPLC for purity check, and identified by LC/MS analysis.

### Structure determination by X-ray crystallography

Lyophilized K48-Ub ^init^ was dissolved in 10 mM HCl to a final concentration of 1 mM. The solution was centrifuged at 15,000 × g for 10 min to remove precipitates, and the resulting supernatant was neutralized by adding a 1/10 volume of 1 M Tris-HCl (pH 8.0). Purified UFD1-UT3 (residues 17–196) was immediately mixed with the neutralized K48-Ub ^init^ at a molar ratio of 1:1.5. The complex was subsequently purified by SEC on HiLoad Superdex 75 column pre-equilibrated with 10 mM Tris-HCl (pH 7.2) containing 50 mM NaCl and 5 mM β-mercaptoethanol to remove unbound monomer. The purified UFD1-UT3–K48Ub ^init^ was concentrated to 11 mg mL^-1^, using an Amicon Ultra-15 10,000 MWCO filter (Millipore), and stored at –80 °C until use.

Initial crystallization screening was performed by the sitting drop vapor diffusion method at 20 °C with a Mosquito liquid-handling robot (TTP Labtech). We tested about 700 conditions using crystallization reagent kits supplied by Hampton Research and Qiagen, and the initial hits were further optimized. The best crystals of the UFD1-UT3–K48Ub_2_^init^ complex were obtained at 20 °C with the sitting drop vapor diffusion method by mixing 1 μL of protein solution with an equal amount of precipitant solution containing 100 mM Bis-Tris (pH 5.5), 200 mM ammonium acetate, and 24% PEG 3350. The drops were equilibrated against 500 μL of the precipitant solution. For data collection, the crystals were transferred to a cryostabilizing solution (the precipitant solution containing 20% ethylene glycol) and flash-frozen in liquid nitrogen.

Diffraction data sets were collected to a maximum resolution of 1.31 Å on beamline BL32XU at SPring-8 (Hyogo, Japan) and processed with XDS^43^ and the CCP4^44^ program suite. The UFD1-UT3–K48Ub_2_ ^init^ structure was determined by molecular replacement, using Molrep^45^. The AF3 model of UFD1-UT3 and the crystal structure of Ub were used as the search models. The atomic models were manually adjusted using Coot^46^ with careful inspection, and refined using Phenix^47^. The final model was obtained after iterative correction and refinement of the models. All molecular graphics were prepared using ChimeraX^42^.

### Preparation of biotinylated K48-Ub_2_

Lyophilized biotinylated Ub was dissolved following the procedure for K48-Ub ^init^. Biotinylated K48-Ub_2_ was synthesized by incubating 300 μM biotinylated Ub and 450 μM Ub (K48R) with 2 μM Ub-activating enzyme E1 and 10 μM E2-25K in a ubiquitination buffer containing 50 mM Tris-HCl (pH 9.0), 10 mM ATP, 10 mM MgCl₂, and 0.6 mM DTT at 37 °C for 18 h. The reaction product was purified using a Resource S cation-exchange column (Cytiva).

### BLI analysis of the interactions between UFD1-UT3 and ubiquitin variants

Affinity measurement was performed with a streptavidin biosensor (ForteBio) using the Octet system (ForteBio) as described in the manufacturer’s instructions. Biotinylated Ub, K48-Ub_2_ and K48-Ub ^init^ was dissolved in HBSTP buffer [50 mM HEPES (pH 7.5), 300 mM NaCl, 0.05% tween 20, 0.1% PEG] and immobilized on the streptavidin sensor. The analyte protein (WT or mutant GST-tagged UFD1) was also dissolved in HBSTP buffer (80 nM final conc.) and binding assay was performed at 25 °C with equilibration for 60 s, association for 600 s, and dissociation for 600 s. For the estimation of dissociation constants, 4 different concentrations of analyte (10, 20, 40, 80 nM) were used to determine kinetic parameters. The obtained data were fitted to a 1:1 binding model using ForteBio Data Analysis 10.0 software.

### Preparation of the polyubiquitinated substrate for unfolding assay

For the polyubiquitination of fluorescence-labeled substrates, 20 μM Ub- or Ub_4_-mEos3.2–His_8_, 1 μM Ub-activating enzyme E1, 20 μM gp78RING–UbE2G2 chimera, and 1000 μM Ub were incubated in ubiquitination buffer at 37 °C for 18 h. To remove free Ub and Ub chains and enzymes, polyubiquitinated Ub- or Ub_4_-mEos3.2-His_8_ was purified using a Ni-NTA column (Qiagen). The eluate was further purified by SEC on Enrich SEC 650 (Bio-Rad) column equilibrated with 50 mM HEPES–NaOH (pH 7.5) containing 100 mM NaCl. Fractions containing Ub- or Ub_4_-mEos3.2-His_8_ modified with long Ub chains (estimated to include more than ten Ub moieties) were collected. The concentration of the polyubiquitinated Ub- or Ub_4_-mEos3.2-His_8_ was determined using a NanoDrop 2000c (Thermo Fisher Scientific) by measuring absorbance at 507 nm, with unmodified Ub– or Ub_4_-mEos3.2-His_8_ as a standard.

To induce photoconversion of mEos3.2, polyubiquitinated Ub- or Ub_4_-mEos3.2-His_8_ (∼10 μM) was placed in a 200 μL PCR tube on an ice bath and irradiated with a 395 nm ultraviolet flashlight (TATTU U3) for 1 h. During irradiation, the sample was gently inverted every 15 min to ensure uniform exposure. The photoconverted polyubiquitinated Ub– and Ub_4_-mEos3.2-His_8_ were then stored at –80 °C as the K48- and the M1/K48-substrates, respectively.

### Single-turnover unfolding assay

For the unfolding assay, photoconverted, K48- or M1/K48-substrate (10 nM) was mixed with 200 nM p97 (monomer basis), 100 nM UFD1-NPL4 heterodimer, and 100 nM additional adaptors in unfolding buffer (50 mM HEPES-NaOH (pH 7.5), containing 100 mM NaCl, 10 mM MgCl_2_, and 0.5 g L^-1^ BSA) in a 384-well black microplate (Greiner). After incubation at 37 °C for 10 min, 2 mM ATP was added, and fluorescence was measured every 30 seconds for 30 min at 37 °C using a Nivo F microplate reader (PerkinElmer) with a 540/30 nm excitation filter, 580/20 nm emission filter, and a D565 dichroic mirror. Fluorescence values were corrected by subtracting the background signal of the unfolding buffer, and data were normalized to the initial reading at 0 min (Extended Data Fig. 4). The relative initial velocity was calculated as the decrease in fluorescence intensity at the 10-minute time point relative to the start of measurement.

### Pexophagy assay

pMXs-Puro/mKeima-SKL and pMXs-Puro/3HA-FAF2 plasmids were described as previously^27^. *FAF2* KO HCT116 cells and *FAF2*/*FIP200* double knockout HCT116 cells were established as previously reported^27^. HCT116 cells transiently or stably expressing YFP, mKeima-SKL, and 3HA-FAF2 in a 6-well plate were resuspended in a sorting buffer (phosphate buffer with 2.5% FBS). Analysis was performed with FACSDiva on a FACSAriaIII cell sorter (BD). mKeima was measured using dual-excitation ratiometric pH measurements at 405 nm (pH 7) and 561 nm (pH 4) with 610/20 nm emission filters.

### CHX chase assay

Human HEK293T cells were obtained from ATCC and maintained at 37°C with 5% CO_2_ in high-glucose Dulbecco’s Modified Eagle’s Medium (DMEM) (Merck) supplemented with 10% fetal bovine serum (FBS) (Merck), penicillin–streptomycin (100 U/mL, GIBCO #15140148), sodium pyruvate (1 mM, GIBCO #11360070), and MEM non-essential amino acids (1×, GIBCO #11140050). Transfections were performed using the Lipofectamine 3000 reagent (Thermo Fisher Scientific) for plasmids and Lipofectamine RNAiMax (Thermo Fisher Scientific) for siRNAs. Transfected cells were used for experiments 48 h (plasmids) or 72 h (siRNA) after transfection. The cells were treated with 100 μg/mL cycloheximide (CHX) for 0, 1, 3, or 6 h before cell lysis. Cells were lysed in a lysis buffer [10 mM Tris-HCl (pH 7.5), 150 mM NaCl, 0.5 mM EDTA, 1% NP-40, and 10% glycerol] supplemented with a protease inhibitor cocktail (Nacalai Tesque) and sonicated extensively using a Handy Sonic (TOMY Seiko). After centrifugation, protein concentrations were determined using a BCA assay kit (Takara).

For immunoblot assays, chemiluminescence signals were detected using a FUSION imaging system (Vilber-Lourmat). The following antibodies were used: p21 (Cell Signaling Technologies, #2947, 1:1000), p27 (Cell Signaling Technologies, #3686, 1:1000), TXNIP (Cell Signaling Technologies, #14715, 1:1000), HIF-1α (Cell Signaling Technologies, #14179, 1:1000), FAF1 (Cell Signaling Technologies, #4932, 1:1000), UBXD7 (Thermo Fisher Scientific, #PA5-71280, 1:500), HA (Cell Signaling Technologies, #3724, 1:1000), and CDT1 (Cell Signaling Technologies, #8064, 1:1000).

### Analysis of p97 complex stoichiometry by SEC

Purified p97 (50 μM), UN (25 μM), and either UBXD7 or FAF1 (50 μM) were mixed in the presence of 1 mM ATP-γS. Following incubation on ice for at least 1 hour, the mixtures were subjected to SEC on a Superdex 200 5/150 GL column (Cytiva) equilibrated with 50 mM Na-HEPES (pH 7.5), 150 mM NaCl, 5 mM MgCl_2_, and 0.5 mM TCEP.

## Extended Data Figure Legends

**Extended Data Figure 1.**
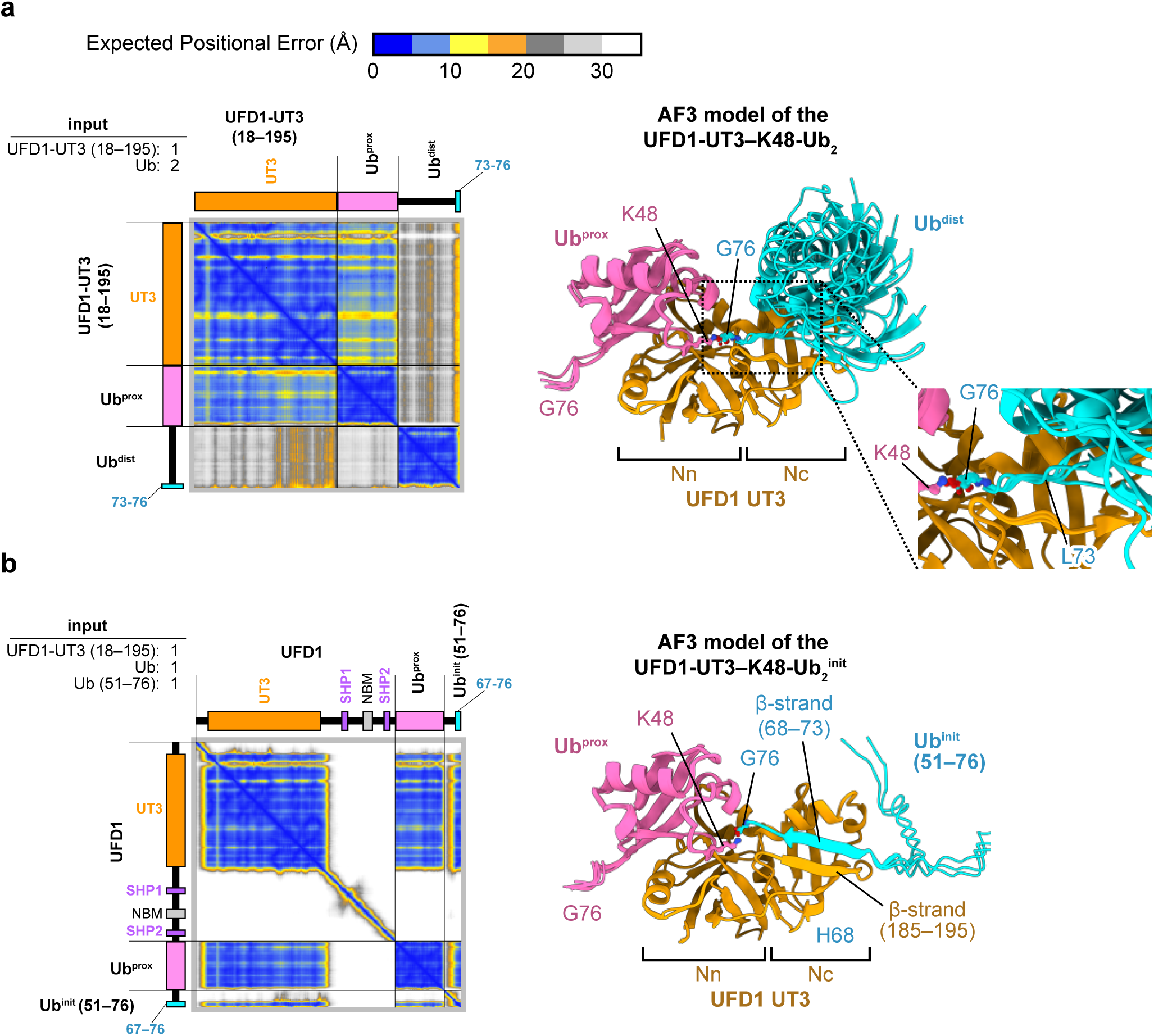
AF3-predicted structures of the UFD1-UT3 in complex with K48-Ub_2_. **a,** PAE plots and predicted structures of the complex generated using one chain of UFD1-UT3 (residues 18–195) and two chains of full-length Ub. Despite the absence of inter-protein distance restraints, the prediction positioned Gly76 of one Ub near Lys48 of the other, resulting in the UFD1-UT3–K48-Ub_2_ complex model. The right panel shows a superposition of the five structures from the AlphaFold Server^28^, aligned on UFD1-UT3. UFD1-UT3, Ub^prox^, and Ub^dist^ are colored orange, pink, and cyan, respectively. **b,** PAE plots and predicted structures generated using one chain each of full-length UFD1, full-length Ub, and a Ub C-terminal fragment (residues 51–76). Despite the absence of inter-protein distance restraints, the prediction positioned Gly76 of the unfolded Ub C-terminal fragment (residues 51–76) near Lys48 of the full-length Ub, yielding the UFD1-UT3–K48-Ub ^init^ complex model. The right panel shows a superposition of five structures aligned on UFD1-UT3. For full-length UFD1, only the UFD1-UT3 region is shown. UFD1-UT3, Ub^prox^, and Ub^init^ are colored orange, pink, and cyan, respectively. In **a** and **b**, protein boundaries in the PAE plots are indicated by black lines. The plots are colored according to the legend, with blue representing high confidence (0–10 Å), yellow and orange indicating moderate confidence (10–20 Å), grey representing low confidence (20–30 Å), and white indicating no confidence (>30 Å). The names of the input proteins and the number of chains are shown in the top left corner of the PAE plot. Corresponding protein names and domain architectures are labeled above and to the left of the plots.

**Extended Data Figure 2.**
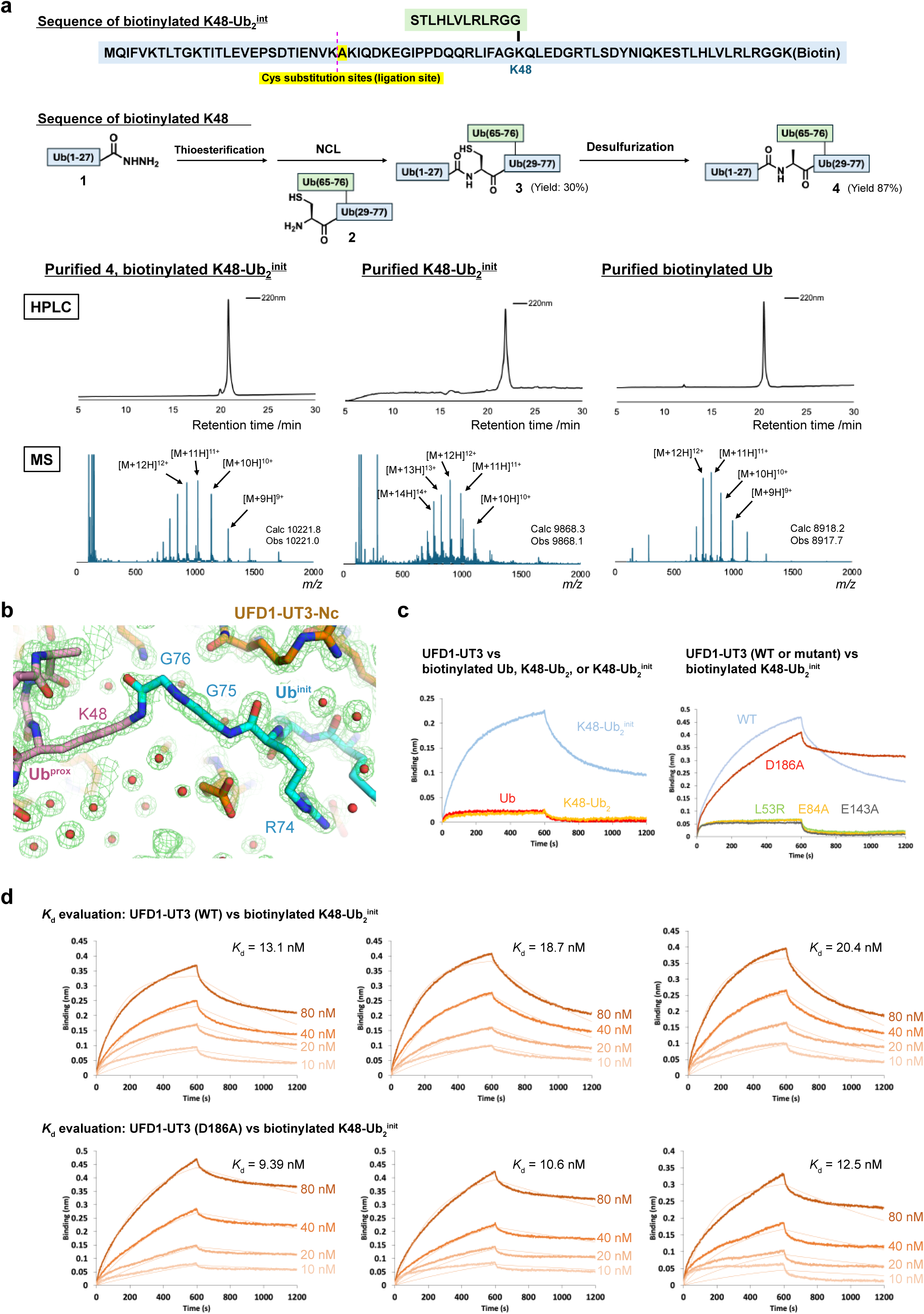
Interaction between UFD1-UT3 and chemically synthesized K48-Ub ^init^. **a,** Chemical synthesis of biotinylated K48-Ub_2_ ^init^, biotinylated Ub and K48-Ub ^init^. The sequence and synthetic scheme of biotinylated K48-Ub ^init^ are shown at the top. Analytical HPLC charts and MS spectra of purified products are shown at the bottom. HPLC peaks were monitored at 220 nm in the linear gradient with water/acetonitrile containing 0.1% TFA. **b,** Close-up view of the area around the isopeptide linkage of chemically synthesized K48-Ub ^init^. Color coding for the protein model follows the scheme in Fig. 1a. A 2*F*_o_–*F*_c_ map is shown as a green mesh contoured at 1.5 σ level. **c, d,** BLI analyses with K48-Ub ^init^ and UFD1. Binding tests with different types of biotinylated Ub derivatives, K48-Ub ^init^, mono-Ub and K48-Ub_2_ (c left) and different types of UFD1 constructs, wildtype and 4 mutants (c right) are shown. Dissociation constant estimation of the K48-Ub2init/WT-UFD1 pair and the K48-Ub ^init^/D186A mutant UFD1 pair are shown in d.

**Extended Data Figure 3.**
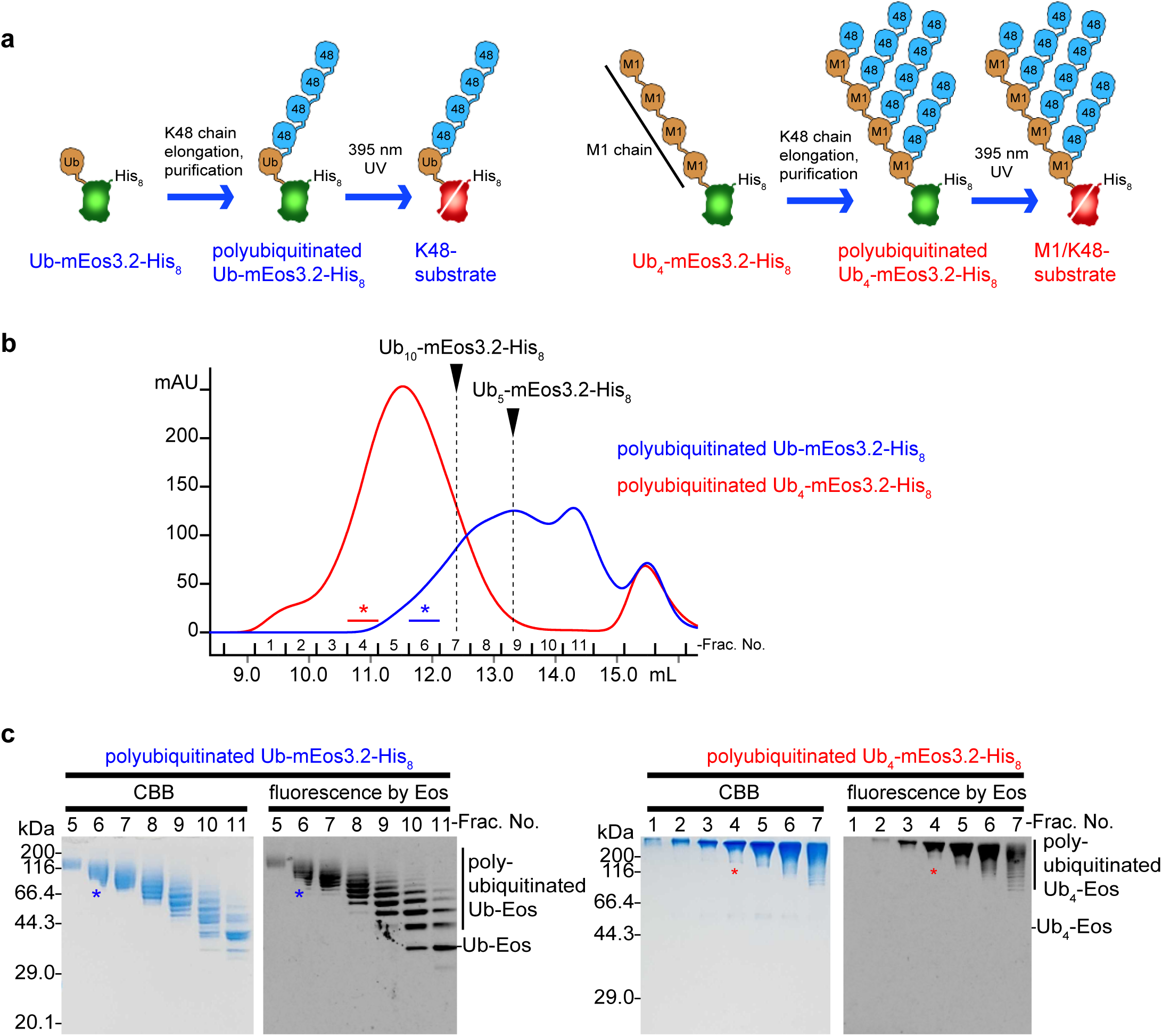
Preparation of K48-chain-modified substrates. **a,** Schematic view of the substrate preparation protocol. The K48-substrate was generated from a fusion protein consisting of mEos3.2 with an N-terminal Ub and a C-terminal His_8_ tag. This construct was enzymatically modified with K48-linked Ub chains by E1 and a gp78RING-Ube2G2 chimera, followed by Ni-NTA and size-exclusion chromatography (SEC) purification. The final product was obtained by UV irradiation (395 nm) on ice (left panel). The M1/K48-substrate was prepared in the same manner as the K48-substrate, except that mEos3.2 fused with an N-terminal M1-Ub_4_ and a C-terminal His_8_ tag was used as the starting material. **b,** SEC chromatograms for polyubiquitinated Ub-mEos3.2-His_8_ (blue line) and Ub_4_-mEos3.2-His_8_ (red line). Absorbance was monitored at 280 nm. Dotted lines indicate positions corresponding to the peaks of mEos3.2-His_8_ modified with K48-Ub_5_ or K48-Ub_10_. Fractions marked with blue (K48-substrate) or red (M1/K48-substrate) asterisks were collected and used for unfolding assays after UV irradiation at 395 nm. **c,** SDS-PAGE analysis of the SEC fractions shown in b. The gel was visualized via Eos-derived fluorescence and subsequently stained with Coomassie Brilliant Blue (CBB).

**Extended Data Figure 4.**
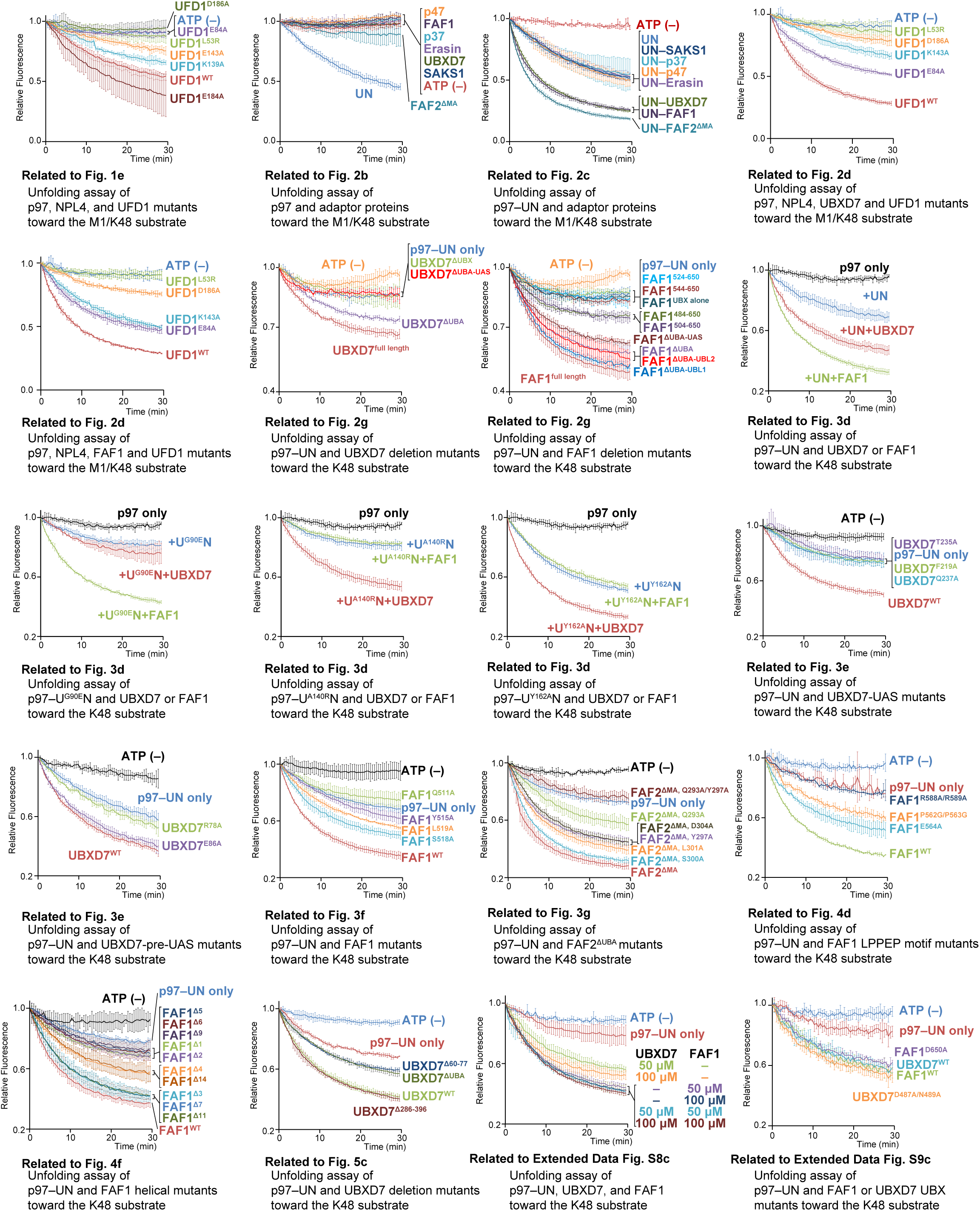
Kinetic traces for p97-mediated substrate unfolding. p97 was incubated with the indicated adaptor proteins and either the K48-substrate or the M1/K48-substrate. Curves represent the mean +/- s.d. (n = 3 independent replicates). These fluorescence decay curves provided the raw data used to calculate the initial velocities shown in the main figures.

**Extended Data Figure 5.**
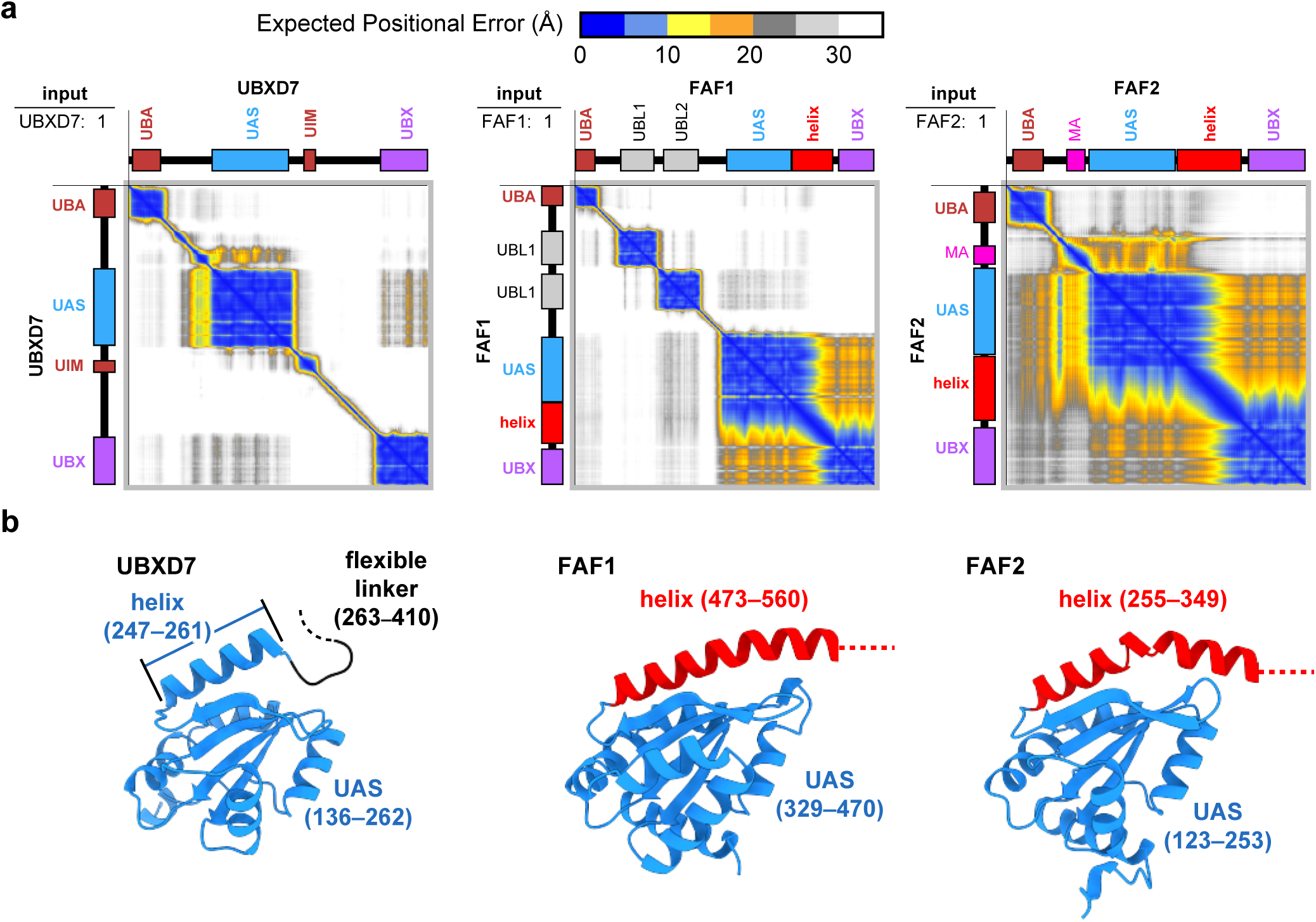
AF3 structural predictions for UBXD7, FAF1, and FAF2. **a,** PAE plots for UBXD7, FAF1, and FAF2 predicted by AF3. The corresponding predicted structures are shown in Fig. 2e. The PAE plot representations follow the style in Extended Data Fig. 1. **b,** Close-up views of the UAS domains and the following helices for UBXD7, FAF1, and FAF2. The UAS domains are oriented in the same direction to facilitate structural comparison.

**Extended Data Figure 6.**
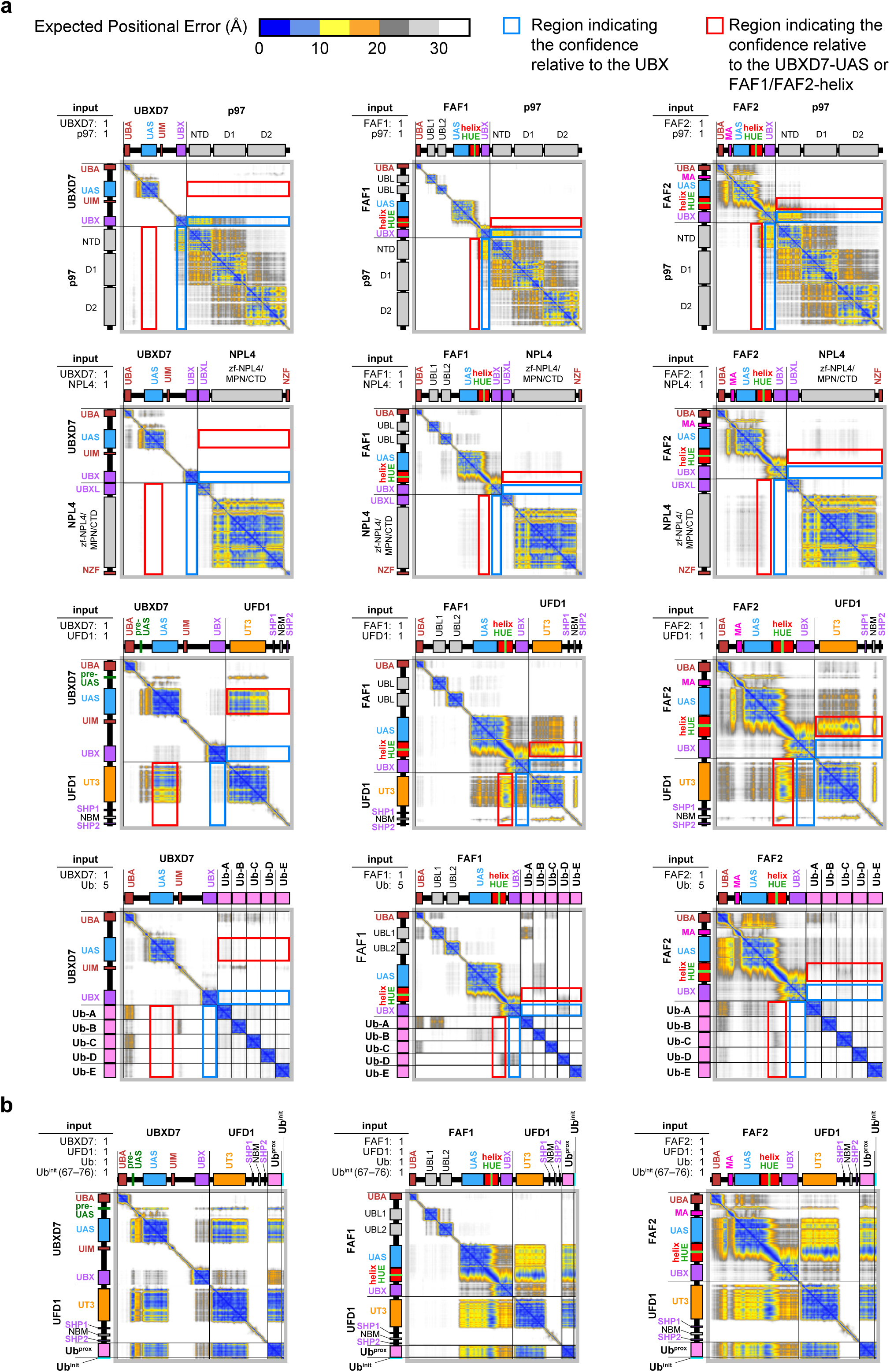
AF3-based screening for interaction partners of UBXD7, FAF1, and FAF2. **a,** PAE plots from AF3 predictions using inputs consisting of a single chain of UBXD7, FAF1, or FAF2 combined with either p97, NPL4, UFD1 (one chain each), or Ub (five chains). Blue and red boxes indicate the regions corresponding to the positional relationships for the UBX domain and the UBXD7-UAS or FAF1/FAF2-HUE motifs, respectively. **b,** PAE plots for interactions between UBXD7, FAF1, or FAF2 and UFD1–K48Ub2^init^. Corresponding predicted structures are shown in Figs 3a–c. In **a** and **b**, the PAE plot representations follow the style in Extended Data Fig. 1. For UBXD7, the position of the pre-UAS region is labeled above and to the left of the plots only when it adopts a structured conformation.

**Extended Data Figure 7.**
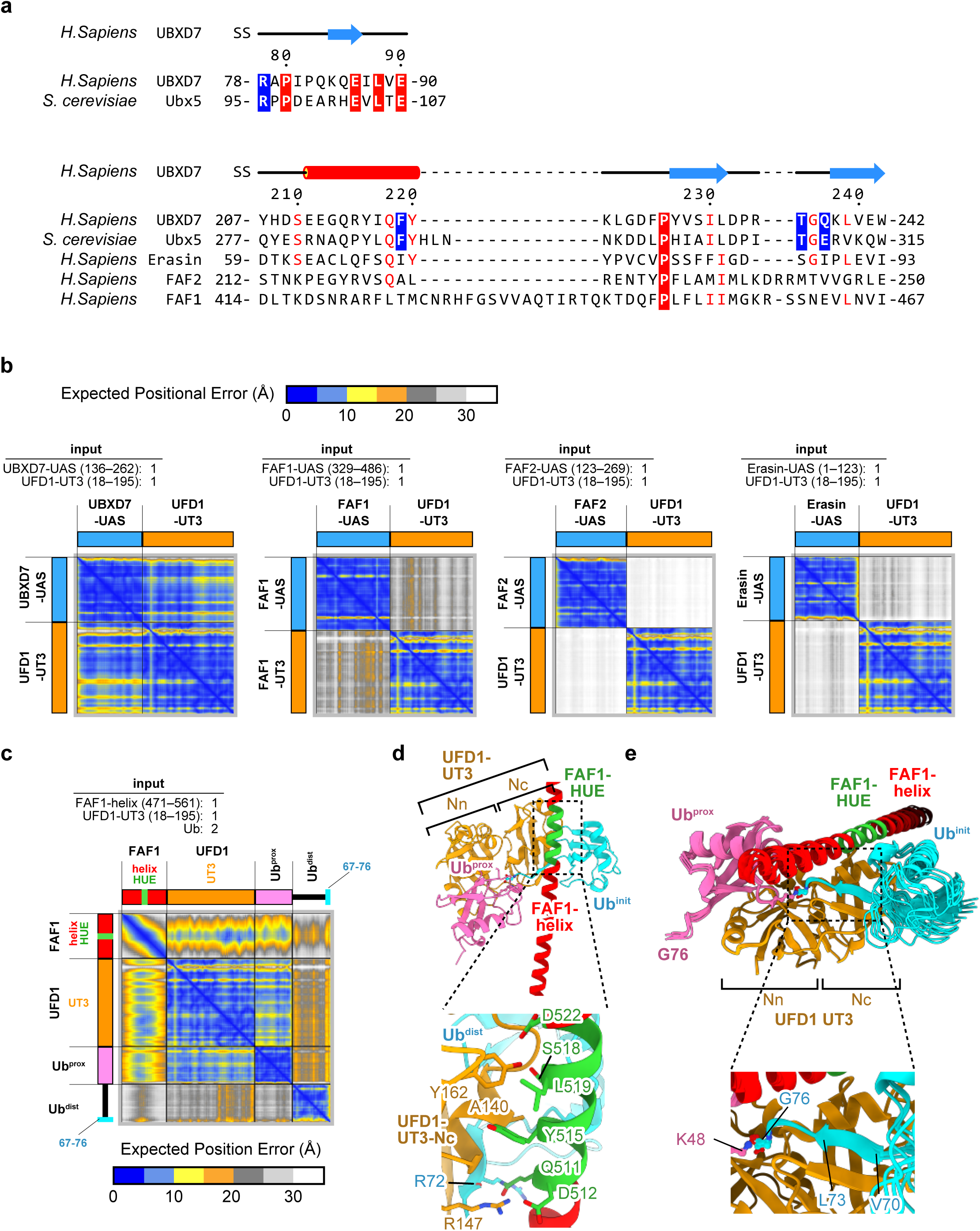
Detailed analysis of UBXD7 and FAF1 interaction with UFD1-UT3. **a,** Sequence alignments of the pre-UAS regions from UBXD7 and its yeast homolog Ubx5 (top), and the UAS domains from UBXD7, Ubx5, Erasin, FAF2, and FAF1 (bottom). Within each alignment, residues identical among all sequences are highlighted in white text on a red background, while those with at least 60% conservation are shown in red text. Residues validated as essential for p97–UN enhancement via UBXD7 mutant analysis are highlighted in white text on a blue background. **b,** PAE plots from AF3 predictions using the UAS domains of UBXD7, FAF1, FAF2, or Erasin with UFD1-UT3. **c, d,** PAE plot (c) and predicted structure (d) of the complex generated using one chain each of FAF1-helix and UFD1-UT3 with two chains of Ub. Despite the absence of inter-protein distance restraints, the prediction positioned Gly76 of one Ub near Lys48 of the other, resulting in the FAF1-helix–UFD1-UT3–K48-Ub_2_ complex model. Components are colored according to Fig. 3b, with Ub^dist^ is colored cyan. In **b** and **c**, the PAE plot representations follow the style in Extended Data Fig. 1. **e,** Superposition of the five FAF1-helix–UFD1-UT3–K48-Ub_2_ complex models obtained from the AlphaFold Server, aligned on UFD1-UT3.

**Extended Data Figure 8.**
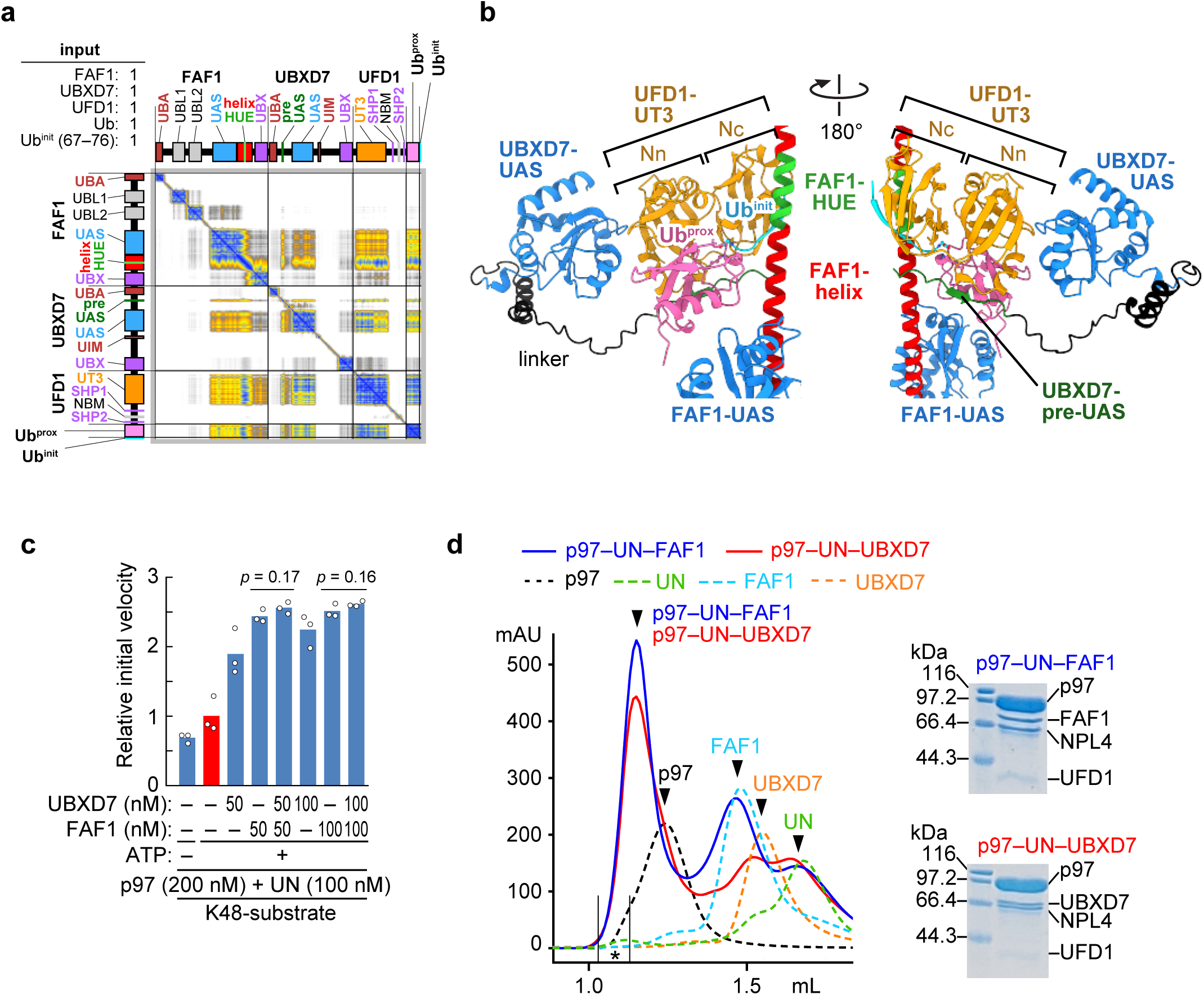
Analysis of the synergistic activation of p97–UN by FAF1 and UBXD7. **a, b,** PAE plot (a) and predicted structure (b) generated from AF3 using FAF1, UBXD7, UFD1, Ub, and the C-terminal Ub^init^ fragment (residues 67–76) as inputs. The PAE plot representations follow the style of Extended Data Fig. S1. Structures are colored according to Fig. 3a and b. Regions not involved in the interactions are omitted for clarity. **c,** Unfolding activity of p97–UN toward the K48-substrate in the presence of FAF1, UBXD7, or both adaptors. Individual data points are shown as white dots, and bars represent the mean (n = 3 independent experiments). Relative initial velocities were normalized to the mean of the p97–UN complex in the presence of ATP (red bar). Statistical significance was determined using one-way ANOVA followed by Dunnett’s *post hoc* tests. Source data are provided in Extended Data Fig. 4. **d,** Purification of p97–UN–FAF1 and p97–UN–UBXD7 complexes by SEC. p97, UN, and either FAF1 or UBXD7 were mixed at a 6:3:6 molar ratio in the presence of 1 mM ATP-γS. SEC analysis showed that both complexes eluted as peaks, with the peaks shifting toward higher molecular weights than those of the individual proteins. SDS-PAGE analysis of the fractions marked with asterisks confirmed that both p97–UN–FAF1 and p97–UN–UBXD7 formed with a 6:1:1 stoichiometry.

**Extended Data Figure 9.**
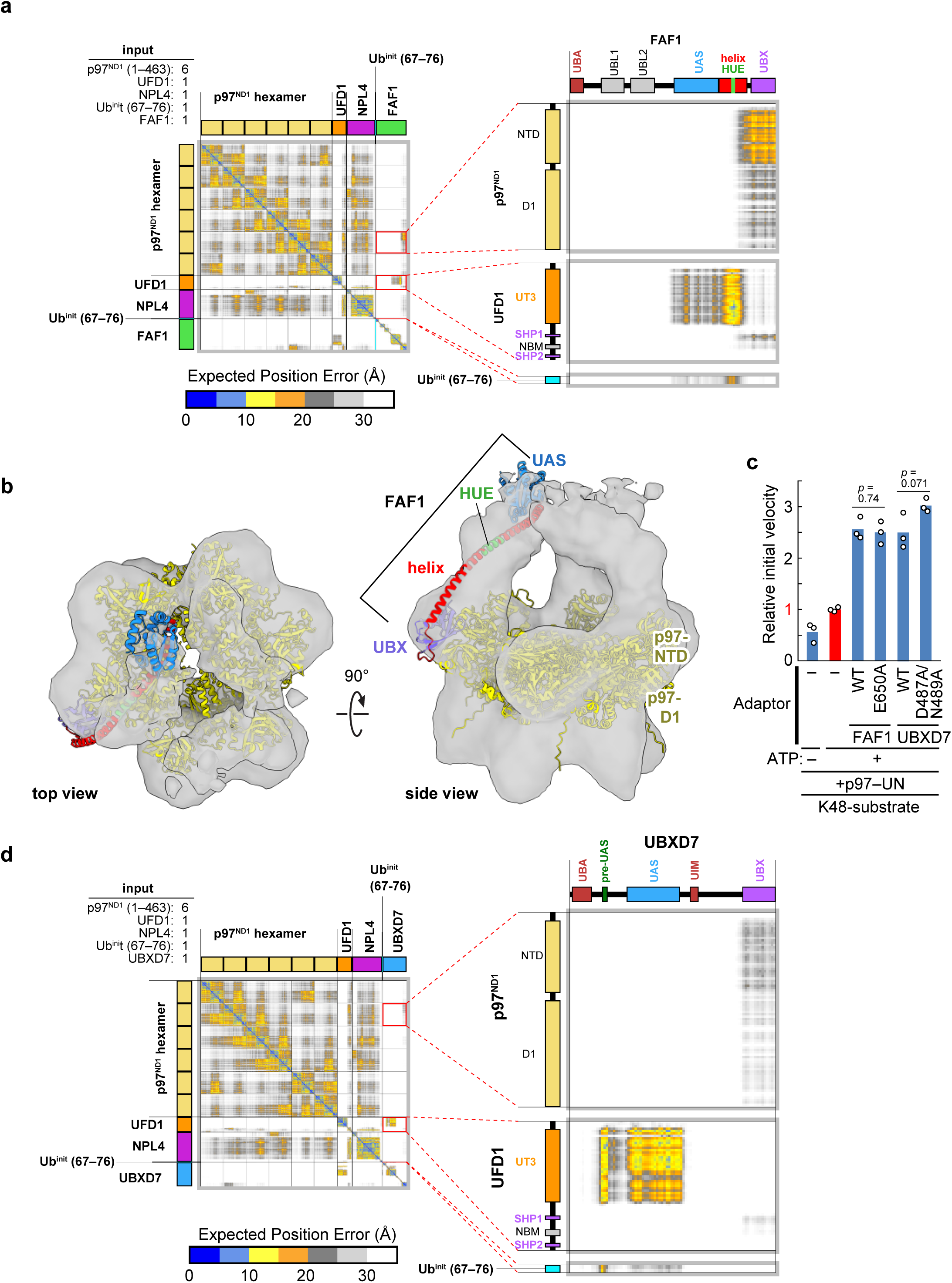
Prediction of the overall structures of the p97–UN complex with FAF1 or UBXD7. **a,** PAE plot for AF3 predictions using six chains of p97^ND1^ (residues 1–463) and one chain each of UFD1, NPL4, FAF1, and the C-terminal Ub^init^ fragment (residues 67–76). The resulting structural model, superimposed with the Ub^init^ fragment (residues 13–50) from the cryo-EM structure (PDB ID: 8DAV)^19^, is presented in Fig. 5a. **b,** Superposition of the isolated p97^ND1^ hexamer and the FAF1 UAS-helix-UBX region from the model in Fig. 4a onto the previously reported EM density map of the 6:3 p97–FAF1 complex (EMD-2319)^35^. Model colors are consistent with those in Fig. 5a. **c,** Impact of mutations in the C-terminal region of the UBX domain of FAF1 and UBXD7 on the unfolding activity of p97–UN toward the K48-substrate. Individual data points are shown as white dots, and bars represent the mean (n = 3 independent experiments). Relative initial velocities were normalized to the mean of the p97–UN complex in the presence of ATP (red bar). Statistical significance was determined using one-way ANOVA followed by Dunnett’s *post hoc* tests. Source data are provided in Extended Data Fig. 4. **d,** PAE plot for AF3 predictions using six chains of p97^ND1^ (residues 1–463) and one chain each of UFD1, NPL4, UBXD7, and the C-terminal Ub^init^ fragment (residues 67–76). The resulting structural model, superimposed with the Ub^init^ fragment (residues 13–50) from the cryo-EM structure (PDB ID: 8DAV)^19^, is presented in Fig. 6a. In **a** and **d**, the PAE plot representations follow the style in Extended Data Fig. 1.

